# DTX3L ubiquitin ligase ubiquitinates single-stranded nucleic acids

**DOI:** 10.1101/2024.04.02.587769

**Authors:** Emily L. Dearlove, Chatrin Chatrin, Lori Buetow, Syed F. Ahmed, Tobias Schmidt, Martin Bushell, Brian O. Smith, Danny T. Huang

**Affiliations:** Cancer Research UK Scotland Institute, Garscube Estate, Switchback Road, Glasgow, G61 1BD, United Kingdom; School of Cancer Sciences, University of Glasgow, Glasgow, G61 1QH, United Kingdom; Sir William Dunn School of Pathology, University of Oxford, Oxford OX1 3RE, United Kingdom; School of Molecular Biosciences, University of Glasgow, Glasgow, G12 8QQ, United Kingdom

## Abstract

Ubiquitination typically involves covalent linking of ubiquitin (Ub) to a lysine residue on a protein substrate. Recently, new facets of this process have emerged, including Ub modification of non-proteinaceous substrates like ADP-ribose by the DELTEX E3 ligase family. Here we show that the DELTEX family member DTX3L expands this non-proteinaceous substrate repertoire to include single-stranded DNA and RNA. Although the N-terminal region of DTX3L contains single-stranded nucleic acid binding domains and motifs, the minimal catalytically competent fragment comprises the C-terminal RING and DTC domains (RD). DTX3L-RD catalyses ubiquitination of the 3’-end of single-stranded DNA and RNA, as well as double-stranded DNA with a 3’ overhang of two or more nucleotides. This modification is reversibly cleaved by deubiquitinases. NMR and biochemical analyses reveal that the DTC domain binds single-stranded DNA and facilitates the catalysis of Ub transfer from RING-bound E2-conjugated Ub. Our study unveils the direct ubiquitination of nucleic acids by DTX3L, laying the groundwork for understanding its functional implications.

## INTRODUCTION

Ubiquitination—the covalent attachment of ubiquitin (Ub) to a substrate—is a versatile post-translational modification involved in the regulation of various cellular functions (*1*). The most widely studied examples of this modification involve conjugation of Ub to lysine residues in proteinaceous substrates. Non-lysine ubiquitination of protein substrates via serine (*2*) and threonine (*3*) residues has also been demonstrated. The modification of these residues produces an ester linkage in contrast to the canonical isopeptide amide bond that is observed in the ubiquitination of lysine residues (*4*). In recent years, non-proteinaceous ubiquitination substrates have emerged. These substrates include bacterial lipopolysaccharide which is ubiquitinated by RNF213 in the immune response to *Salmonella* (*5*), glucosaccharides which are ubiquitinated by HOIL-1 (*6*) and adenosine 5′-diphosphate (ADP)–ribose (ADPr) which is ubiquitinated by the Deltex family of ubiquitin ligases (E3s) (*7–9*). Collectively, these studies have uncovered that the diversity of substrates undergoing ubiquitination expands beyond that of proteinaceous substrates. Additionally, they highlight that this mechanism is not limited to one E3 ligase family.

The Deltex (DTX) E3 family is comprised of five RING E3s, DTX1, DTX2, DTX3, DTX3L and DTX4, which share a conserved C-terminal region containing the RING and DELTEX C-terminal (DTC) domains (*7, 10*). Whilst the RING domain functions by recruiting E2 thioesterified with Ub (E2∼Ub) to catalyse Ub transfer to a substrate (*11*), the DTC domain, which is connected to the RING domain through a short flexible linker remained, until recently, without a reported function. DTX3L, which forms a heterodimer with poly(ADP-ribose) polymerase 9 (PARP9) (*12*), was initially proposed to be an activator of PARP9 ADP-ribosyltransferase activity, with the complex being shown to catalyse ADP-ribosylation of Ub in the presence of nicotinamide adenine dinucleotide (NAD^+^) and ubiquitination components including E1, E2, Ub, Mg^2+^ and ATP (*8*). It was originally proposed that PARP9 was responsible for this activity, despite previous reports of its catalytic inactivity (*13*). When we tested DTX3L independently of PARP9, it was sufficient to catalyse ADP-ribosylation of the C-terminus of Ub(*7*). Notably, the C-terminal RING and DTC domains (hereafter referred to as RD domains) are the minimal fragment required to catalyse this reaction and all members of the DTX family share this capability (*7–9*). Structural analyses of RD domains revealed that the DTC domain contains a pocket that binds ADPr and NAD^+^ (*7, 10*). This enables RD domains to recruit ADPr-modified substrates and catalyse their ubiquitination in a poly-ADP-ribose (PAR)-dependent manner (*10*). In addition, in the presence of NAD^+^ and ubiquitination components, RD domains catalyse ADP-ribosylation of Ub (*7*). The chemical linkage of ADP-ribosylated Ub was recently discovered to occur between the carboxylate group of Ub Gly^76^ and the 3’-hydroxyl group of the adenosine-proximal ribose of ADPr or NAD^+^, highlighting Ub modification of ADPr or NAD^+^ rather than the canonical ADP-ribosylation of substrate that involves the release of nicotinamide from NAD^+^ followed by attachment of ADPr via the C1” atom of the nicotinamide ribose (*9*). Furthermore, recent studies have demonstrated that the RD domains of both DTX2 and DTX3L can ubiquitinate the 3’-hydroxyl group of the adenosine-proximal ribose of free ADPr as well as ADPr moieties present on ADP-ribosylated proteins (*9*) and ADP-ribosylated nucleic acids (*14*).

DTX3L has a unique N-terminus lacking the WWE domains and proline-rich regions found in the other DTX family members. The development of AlphaFold Protein Structure Database (*15*) has allowed prediction of unsolved protein structures to a higher degree of confidence than previously possible. For DTX3L, this allowed the generation of a model of the protein that revealed putative domains in the N-terminal region. When queried in the DALI server, which enables protein structures to be compared in 3D, these domains were found to be structurally similar to K Homology (KH) domains and RNA recognition motifs (RRMs) which bind single-stranded DNA (ssDNA) and RNA (ssRNA). Here, we showed that DTX3L was able to bind a variety of sequences of ssDNA and ssRNA. Unexpectedly, we discovered that DTX3L was able to ubiquitinate ssDNA and ssRNA. Biochemical and structural analyses established that DTX3L-RD domain binds single-stranded nucleic acids (ssNAs) and catalyzes the ubiquitination of ssNAs. These findings present a previously unidentified direct Ub modification of nucleic acids.

## RESULTS

### DTX3L binds and ubiquitinates ssRNA and ssDNA

To gain insight into the function of DTX3L, we took the AlphaFold-predicted structural domains of DTX3L (Fig. 1A) and conducted a search on the DALI server to compare the proposed 3D domains of DTX3L to structures in the Protein Data Bank (PDB). Several of these domains share structural similarity to KH domains (Fig. 1, B and C) with the N-terminal domain resembling an RRM. KH domains, which bind both ssRNA (*16*) and ssDNA (*17*), comprise a three-stranded anti-parallel β-sheet packed alongside three α-helices whereas RRMs are composed of two α-helices and four anti-parallel strands packed together in a β-α-β-β-α-β fold (*18*). To determine if DTX3L binds ssRNA or ssDNA, we used ssDNA and ssRNA oligonucleotides labelled with 6-FAM on the 5’-end (D1-D9 and R1-R9, respectively; Table 1 and fig. S1A) and either full-length (FL) DTX3L or a truncated 232-C construct containing the two tandem KH-like (KHL) domains and C-terminal RING-DTC that is competent in binding PARP9 (*19*). Both DTX3L constructs bound to all tested sequences of ssRNA and ssDNA, inducing at least a two-fold change in polarisation (Fig. 1, D and E and fig. S1, B and C). Likewise, at least a two-fold change in polarisation was observed when DTX3L-PARP9 complex binding was tested with these nucleic acids, demonstrating that PARP9 does not occlude ssDNA or ssRNA binding by DTX3L (fig. S1, D and E).

**Figure 1.**
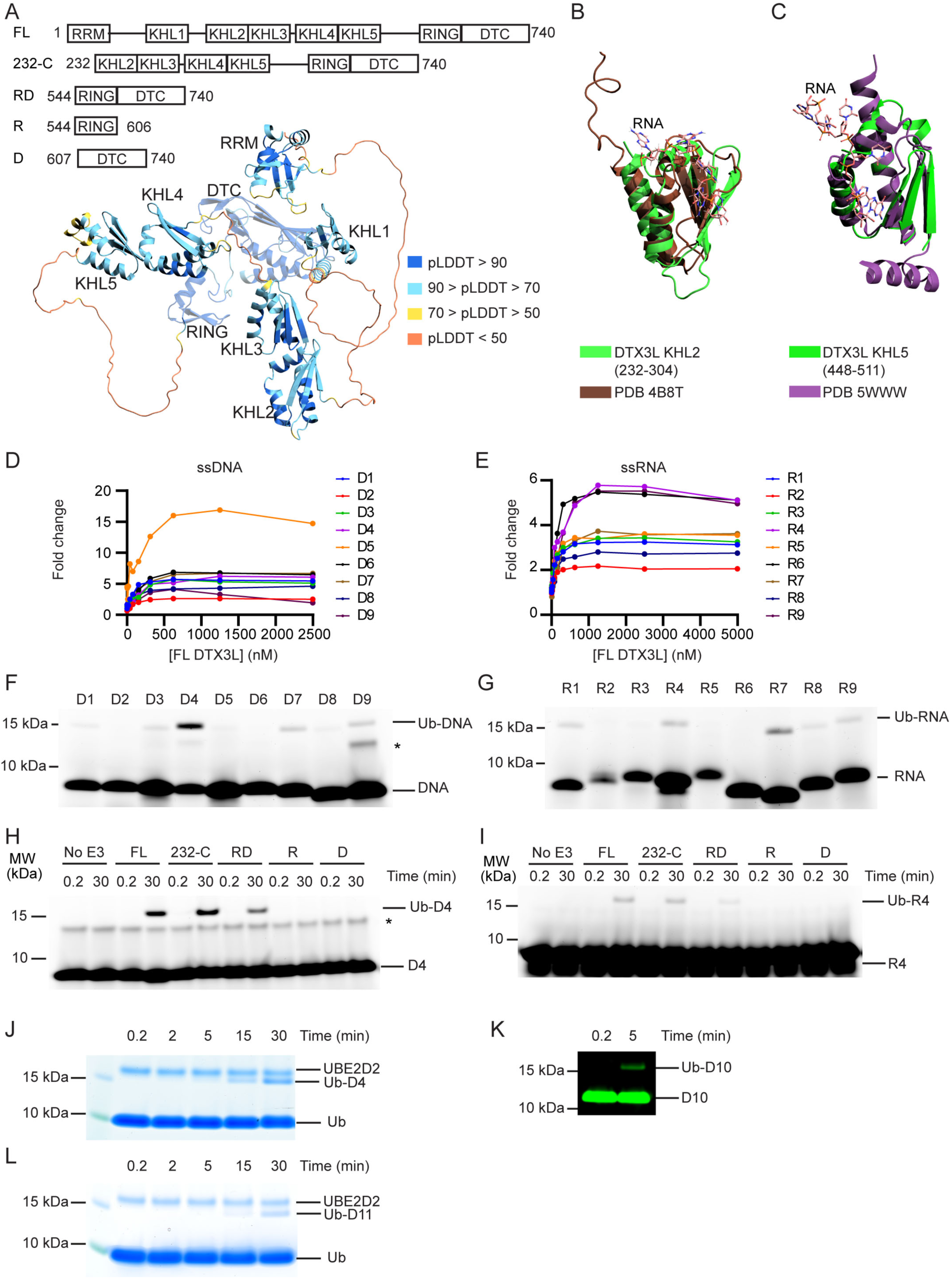
DTX3L catalyses ubiquitination of single-stranded nucleic acids. **(A)** Cartoon representation of the AlphaFold model of DTX3L. Domains are coloured according to model confidence. Domain architecture of DTX3L constructs is shown above the model. **(B)** DTX3L KHL2 (232-304) prediction shown in green overlaid with the third KH domain of KSRP in complex with AGGGU RNA sequence (PDB 4B8T) shown in brown. **(C)** DTX3L KHL5 (448-511) prediction shown in green overlaid with the MEX-3C KH1 domain in complex with GUUUAG RNA sequence (PDB 5WWW) shown in purple. **(D)** Fold change of fluorescence polarisation of 6-FAM-labelled ssDNA D1-9 upon titrating with full-length DTX3L. **(E)** As in (D) but with 6-FAM-labelled ssRNA R1-9. **(F)** Fluorescently detected SDS-PAGE gel of *in vitro* ubiquitination of 6-FAM-labelled ssDNA D1-9 oligonucleotides by FL DTX3L in the presence of E1, UBE2D2, Ub, Mg^2+^-ATP. **(G)** As in (F) but with 6-FAM-labelled ssRNA R1-9 oligonucleotides. **(H)** Fluorescently detected SDS-PAGE gel of *in vitro* ubiquitination of 6-FAM-labelled ssDNA D4 by DTX3L variants (FL, full length; RD, RING-DTC domains; R, RING domain; D, DTC domain) in the presence of E1, UBE2D2, Ub, Mg^2+^-ATP. **(I)** As in (H) but with 6-FAM-labelled ssRNA R4. **(J)** Coomassie stained SDS-PAGE gel of *in vitro* ubiquitination of 6-FAM-labelled ssDNA D4 by DTX3L-RD in the presence of E1, UBE2D2, Ub, Mg^2+^-ATP. (**K**) Fluorescently detected SDS-PAGE gel of *in vitro* ubiquitination of 5’ IRDye® 800 ssDNA D10 by DTX3L-RD in the presence of E1, UBE2D2, Ub, Mg^2+^-ATP. (**L**) Coomassie stained SDS-PAGE gel of *in vitro* ubiquitination of ssDNA D11 by DTX3L-RD in the presence of E1, UBE2D2, Ub, Mg^2+^-ATP. Asterisks in (F) and (H) indicate contaminant bands from ssDNA or ssRNA. Uncropped gel images for (F–L) are shown in fig. S4.

**Table 1.**
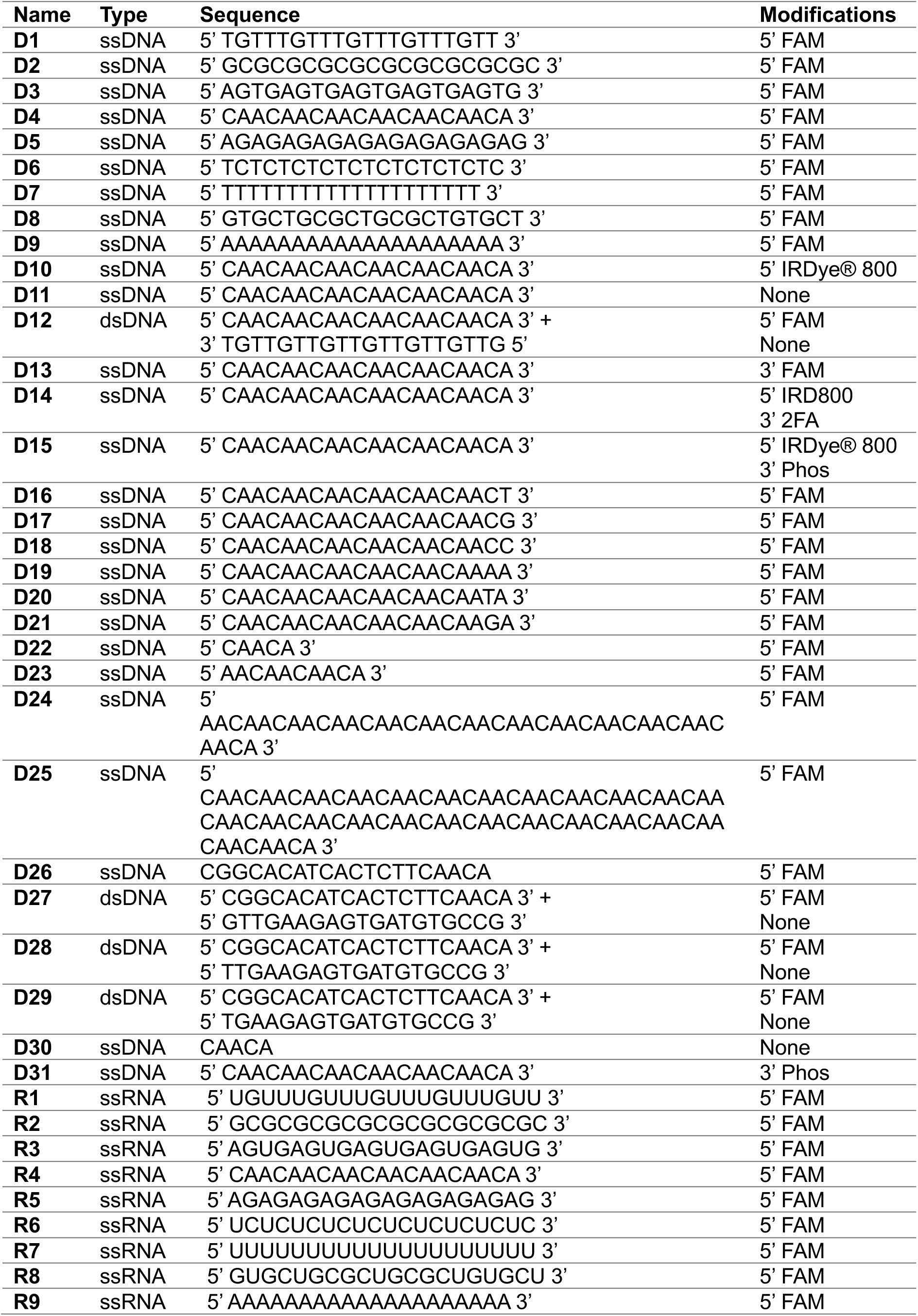
List of nucleotide sequences used in this study.

Based on the structural similarity between ADPr and nucleic acids (fig. S1F), we hypothesised that RNA and DNA could also be ubiquitinated by DTX3L. To investigate, we performed *in vitro* ubiquitination assays of FAM-labelled ssDNA or ssRNA by DTX3L with the following components: E1, UBE2D2, Ub, and Mg^2+^-ATP. Ubiquitination of FAM-labelled ssDNA and ssRNA was detected by SDS-PAGE using a fluorescent scanner (Typhoon FLA7000). The appearance of a ∼15 kDa band closely matched the combined molecular weights of Ub (8564 Da) and 20 nt FAM-labelled single-stranded nucleic acid (∼6900-7300 Da), suggesting that both ssDNA (Fig. 1F) and ssRNA (Fig. 1G) were ubiquitinated by DTX3L. Due to the high signal-to-noise ratio observed with D4 compared to other oligonucleotides, this sequence was used for subsequent experiments. Because R4 is the corresponding sequence of ssRNA this was also taken forward.

The minimal fragment of DTX3L required for Ub-ADPr formation is the conserved C-terminal RING-DTC domain (*7*). To investigate the minimal fragment of DTX3L necessary for the ubiquitination of ssNAs, we tracked the formation of ubiquitinated ssDNA (Ub-DNA) or ssRNA (Ub-RNA) by DTX3L 232-C, RD, RING or DTC. DTX3L 232-C lacks the N-terminal RRM and KHL1 domains, where KHL1 domain was recently proposed to enable oligomerization (*20*). While DTX3L 232-C exhibited reduced autoubiquitination compared to DTX3L FL (fig. S1G), the rates of UBE2D2∼Ub discharge were comparable (fig. S1H). This suggests that DTX3L 232-C is catalytically competent as DTX3L FL, but lacks accessible lysine sites necessary for autoubiquitination. DTX3L 232-C exhibited similar activity as DTX3L FL in catalysing the formation of Ub-DNA (Fig. 1H) and Ub-RNA (Fig. 1I). DTX3L-RD also catalysed formation of Ub-DNA (Fig. 1H) and Ub-RNA (Fig. 1I), whereas neither the RING (R) nor the DTC (D) alone sufficed. Hints of the ∼15 kDa Ub-DNA band were also observed on the corresponding Coomassie-stained SDS-PAGE gel (fig. S1I, arrows).

We took advantage of the Coomassie staining method of detection for Ub-DNA, along with an increased concentration of D4, to confirm this and demonstrate that formation of Ub-DNA is time-dependent (Fig. 1J). To validate that Ub modification did not occur on the FAM label, we tested ssDNA labelled with IRDye® 800 (D10, Table 1; fig. S1J) and unlabelled ssDNA (D11, Table 1) in a ubiquitination assay. A ∼15 kDa band corresponding to the predicted molecular weight of Ub-DNA was also observed using ssDNA with the alternative label (Fig. 1K) and unlabelled ssDNA (Fig. 1L). Together, our data demonstrate that DTX3L binds and ubiquitinates ssNAs and that DTX3L-RD is the minimal fragment required to catalyse this reaction.

### The 3’-end of ssDNA is modified on its 3’ hydroxyl

Based on the finding that the RD was sufficient to catalyse ubiquitination of ssDNA D4, we confirmed that this fragment was able to ubiquitinate the same ssDNAs as FL DTX3L (fig. S1K). Hence, this fragment was used to probe the additional requirements of the reaction. Initially, to confirm that product formation was dependent on the ubiquitination process, we removed each reactant individually. Formation of Ub-DNA or Ub-RNA by DTX3L-RD was only detected when all components of the ubiquitination cascade (E1, UBE2D2, Ub, Mg^2+^-ATP) were present (Fig. 2, A and B). To validate that ssDNA and ssRNA were modified with Ub and not another component in the reaction, we treated the resulting product with USP2, a promiscuous deubiquitinating enzyme. The ∼15 kDa band disappeared upon treatment with USP2, which is consistent with removal of Ub from ssDNA (Fig. 2C) and ssRNA (Fig. 2D) and shows that this modification is reversible. Treatment of Ub-DNA and Ub-RNA with Benzonase, an endonuclease which degrades DNA and RNA, caused the disappearance of both the Ub-DNA and DNA bands (Fig. 2C) and Ub-RNA and RNA bands (Fig. 2D). Poly(ADP-ribose) glycohydrolase (PARG), an ADPr hydrolase that cleaves ADPr-linked to substrate, was recently shown to cleave Ub-ADPr attached to nucleic acid substrates (*14*). Treatment with PARG had no effect on the Ub-DNA band (Fig. 2C) or Ub-RNA band (Fig. 2D), thereby demonstrating that this reaction is distinct from the attachment of Ub to the ADPr moiety on ADP-ribosylated nucleic acids (*14*). These findings showed that the 15 kDa product was ubiquitinated ssNAs.

**Figure 2.**
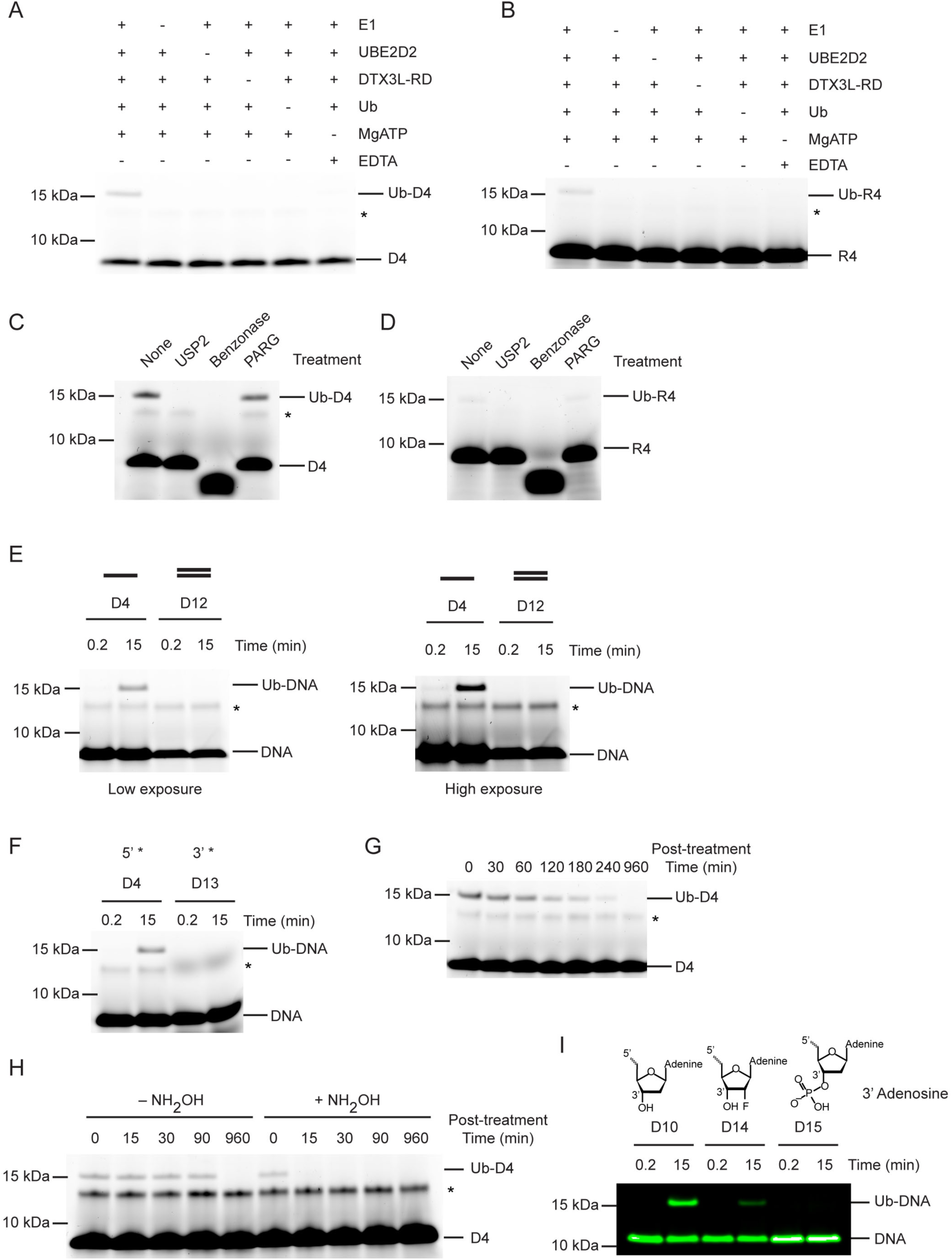
Ub modification of ssDNA occurs at the 3’ hydroxyl at the 3’ end. **(A)** Fluorescently detected SDS-PAGE gel of *in vitro* ubiquitination of ssDNA D4 in which E1, UBE2D2, DTX3L-RD, Ub or Mg^2+^-ATP have been omitted. **(B)** As in (A) but with 6-FAM-labelled ssRNA R4. **(C)** Fluorescently detected SDS-PAGE gel of i*n vitro* ubiquitination of 6-FAM-labelled ssDNA D4 by DTX3L-RD in the presence of E1, UBE2D2, Ub, Mg^2+^-ATP subsequently treated with USP2, Benzonase, PARG or not treated (None). **(D)** As in (C) but with 6-FAM-labelled ssRNA R4. **(E)** Fluorescently detected SDS-PAGE gel of *in vitro* ubiquitination of 6-FAM-labelled ssDNA D4 and dsDNA D12 oligonucleotides by DTX3L-RD (left panel) and at increased exposure (right panel) in the presence of E1, UBE2D2, Ub, Mg^2+^-ATP. **(F)** As in (E) but with 6-FAM-labelled ssDNA D4 (5’ label) and D13 (3’ label) oligonucleotides. **(G)** Fluorescently detected SDS-PAGE gel of *in vitro* ubiquitinated of 6-FAM-labelled ssDNA D4 subsequently treated with pH 9.5 buffer for the times indicated. **(H)** Fluorescently detected SDS-PAGE gel of *in vitro* ubiquitinated of 6-FAM-labelled ssDNA D4 subsequently treated with 1.5 M NH_2_OH at pH 9 for the times indicated. **(I)** As in (E) but with 5’ IRDye® 800 ssDNA D10, D14 and D15 oligonucleotides. Asterisks in (A–C) and (E–H) indicate contaminant band from ssDNA or ssRNA. Uncropped gel images are shown in fig. S4.

Our experiments suggested a common mechanism of modification between ssRNA and ssDNA, therefore we focused our subsequent experiments on the most readily detectable substrate, ssDNA D4. To ascertain whether this reaction was specific to ssDNA, we followed the formation of Ub-DNA with 6-FAM-labelled dsDNA and DTX3L-RD. Based on the finding that the RD fragment was sufficient to catalyse ubiquitination of ssDNA, this fragment was used to probe the additional requirements of the reaction. DTX3L-RD did not ubiquitinate dsDNA (D12, Table 1; Fig. 2E) suggesting that the protein is specific for ssDNA. We then compared ubiquitination of ssDNA labelled with 6-FAM on the 5’-end or 3’-end (D13, Table 1; fig. S1, A and L) and found that the 6-FAM label on the 3’-end blocked ubiquitination of ssDNA (Fig. 2F). Based on this finding we hypothesised that the 3’ end of the oligonucleotide is (1) the site of modification or (2) important for oligonucleotide binding. We next investigated the chemical nature of the covalent bond between Ub and ssDNA. To see if the bond was hydrolysed under basic conditions, we treated our reaction with pH 9.5 buffer which led to the disappearance of the Ub-DNA band over time (Fig. 2G). Additionally, treatment with NH_2_OH resulted in the rapid disappearance of the Ub-DNA band (Fig. 2H). Together, this suggested that the linkage is likely to be an ester, and that the modification occurs on a ribose moiety, leaving only the 3’ ribose as a viable modification site. To validate this hypothesis, we designed an array of 5’-FAM labelled ssDNAs in which various positions on the ribose ring of the 3’-end nucleotide were modified and tested their ability to act as a substrate for ubiquitination by DTX3L-RD. Whilst ssDNA with a fluorine atom attached at the 2’ position of the ribose ring (D14) was ubiquitinated, the addition of a phosphate moiety to the 3’ hydroxyl of the ribose ring (D15) ablated the formation of Ub-DNA, thereby confirming this is the site of modification (D10, D14-D15, Table 1; Fig. 2I). This is consistent with the site of Ub modification of ADPr where Ub is also attached to the 3’ hydroxyl of the adenine-proximal ribose (*9*). These data support that Ub modification of ssDNA occurs at the 3’ hydroxyl of the 3’ nucleotide.

### Nucleotide sequence requirements for Ub-DNA formation

How the sequence and length of ssDNA affect Ub-DNA formation is unclear. To investigate, we first probed the base preference at the last or penultimate position by varying the nucleic acid to A, T, C or G of oligonucleotide D4. Whilst only the sequences ending with A and to some extent with G could be ubiquitinated by DTX3L-RD, the requirement for the penultimate nucleotide was less strict, with sequences ending CA, TA and AA all being ubiquitinated (D16-D21, Table 1; Fig. 3, A and B). Thus far we had only tested sequences of 20 nucleotides in length; therefore, we next took the D4 sequence and generated four ssDNA nucleotides with this same CAA repeat unit and ranging in length from five to 80 nucleotides (D22-D25, Table 1). When tested in our assays, all four of these nucleotide substrates were ubiquitinated (Fig. 3, C and D). Despite dsDNA being unable to act as a substrate for ubiquitination, our discovery that ubiquitination occurs on the 3’ nucleotide of ssDNA prompted us to test the hypothesis that dsDNA with a 3’ overhang could be ubiquitinated. Whilst dsDNA with a single 3’ nucleotide overhang could not be ubiquitinated, dsDNA with a two or three 3’ nucleotide overhang could be modified with Ub (D26-D29, Table 1; Fig. 3E). Our data show that DTX3L modification of DNA requires that the two nucleotides at the 3’ end be single-stranded. Overall, these data demonstrate the ability of DTX3L-RD to ubiquitinate ssDNA of varying lengths and specific 3’ dinucleotides.

**Figure 3.**
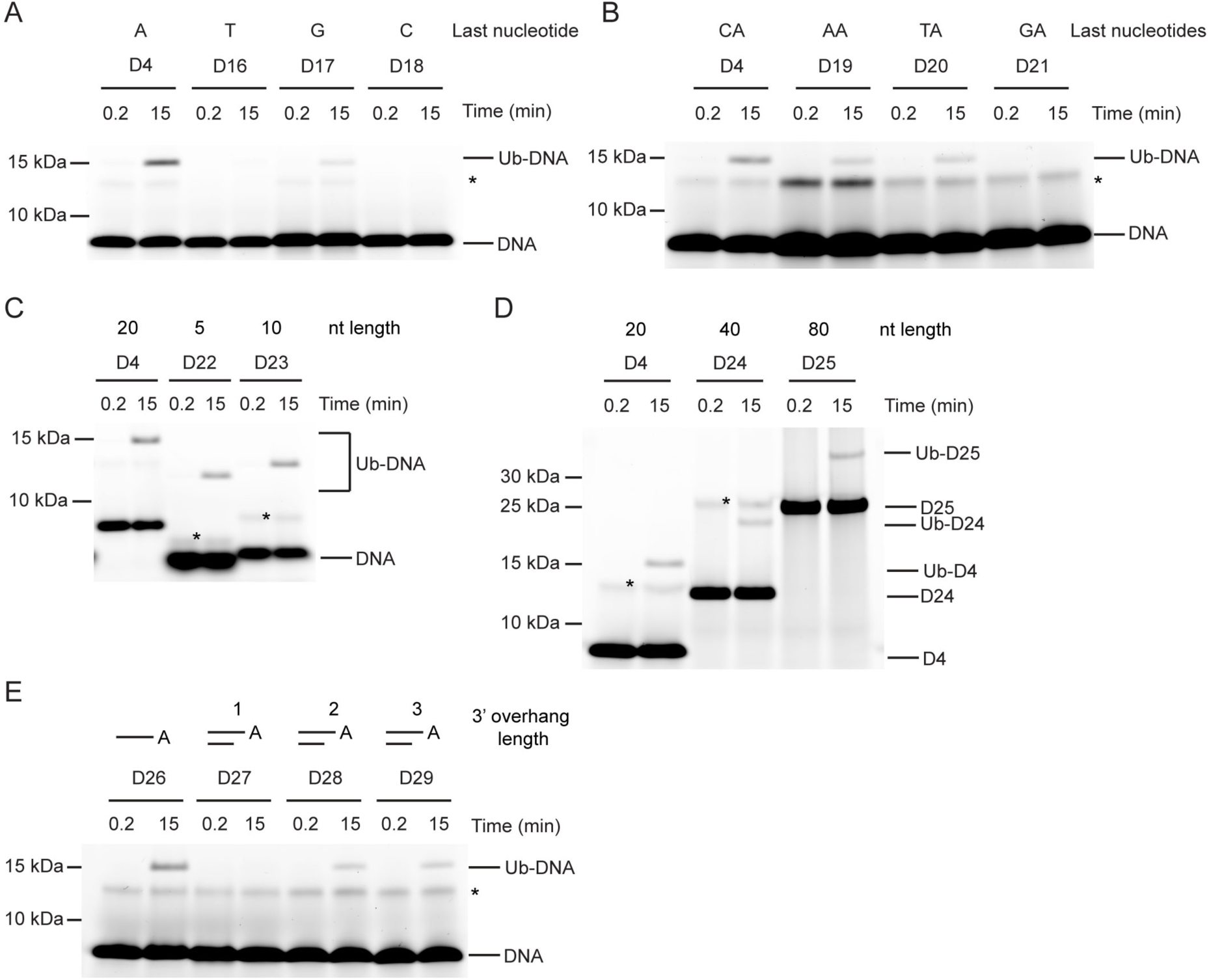
Nucleotide sequence requirements for Ub-DNA formation. Fluorescently detected SDS-PAGE gel of *in vitro* ubiquitination (in the presence of E1, UBE2D2, Ub, and Mg^2+^-ATP) of **(A)** 6-FAM-labelled ssDNA D4, D16, D17 and D18 by DTX3L-RD. **(B)** 6-FAM-labelled ssDNA D4, D19, D20 and D21 by DTX3L-RD. **(C)** 6-FAM-labelled ssDNA D4, D22 and D23 by DTX3L-RD. **(D)** 6-FAM-labelled ssDNA D4, D24 and D25 by DTX3L-RD. **(E)** 6-FAM-labelled ssDNA D26, dsDNA D27, D28 and D29 by DTX3L-RD. Asterisks indicate contaminant band from ssDNA. Uncropped gel images are shown in fig. S4.

### ssNA and ADPr share a binding site on DTX3L’s DTC domain

Because the modification of ssDNA and ADPr occurs at the same position on the nucleoside, we next explored the possibility that they share the same binding site on DTX3L-RD. ADPr binds the DTC domain of DTX2 (*10*), a region which is conserved in the DTX family. Therefore, we purified ^15^N-DTX3L-RD and acquired ^1^H-^15^N heteronuclear single-quantum coherence (HSQC) spectra of ^15^N-DTX3L-RD alone and in the presence of ADPr or unlabelled ssDNA. We utilised the unlabelled version of the 5-mer oligonucleotide ssDNA (D30, Table 1) for NMR analysis, which was sufficient to produce the Ub-DNA product (Fig. 3C). Upon titration of ADPr or 5-nucleotide ssDNA, chemical shift perturbations (CSPs) occurred, indicating DTX3L-RD binds both ADPr and ssDNA (Fig. 4A and fig. S2, A and B). The similarity in the CSPs indicates that there is some overlap in the binding sites of ADPr and ssDNA (Fig. 4, A–C). To further verify that there is conservation between the binding sites of these two substrates, excess ADPr was added to the ubiquitination reaction components along with DTX3L-RD and ssDNA D4. The presence of ADPr inhibited the formation of Ub-DNA and Ub-RNA (Fig. 4D and fig. S2C), suggesting they share the same binding site. We next added excess ssDNA D31, a ssDNA with the same sequence as D4 but with a phosphate moiety attached to the 3’ hydroxyl, to the ubiquitination reaction components along with DTX3L-RD and biotin-NAD^+^. As labelled ADPr is not readily available, we used biotin-NAD^+^ to monitor Ub-biotin-NAD^+^ formation. The addition of D31 inhibited the formation of Ub-NAD^+^ (Fig. 4E). Because D31 has a phosphate moiety blocking the 3’ hydroxyl ubiquitination site, it cannot be ubiquitinated; therefore, it must be directly competing for binding.

**Figure 4.**
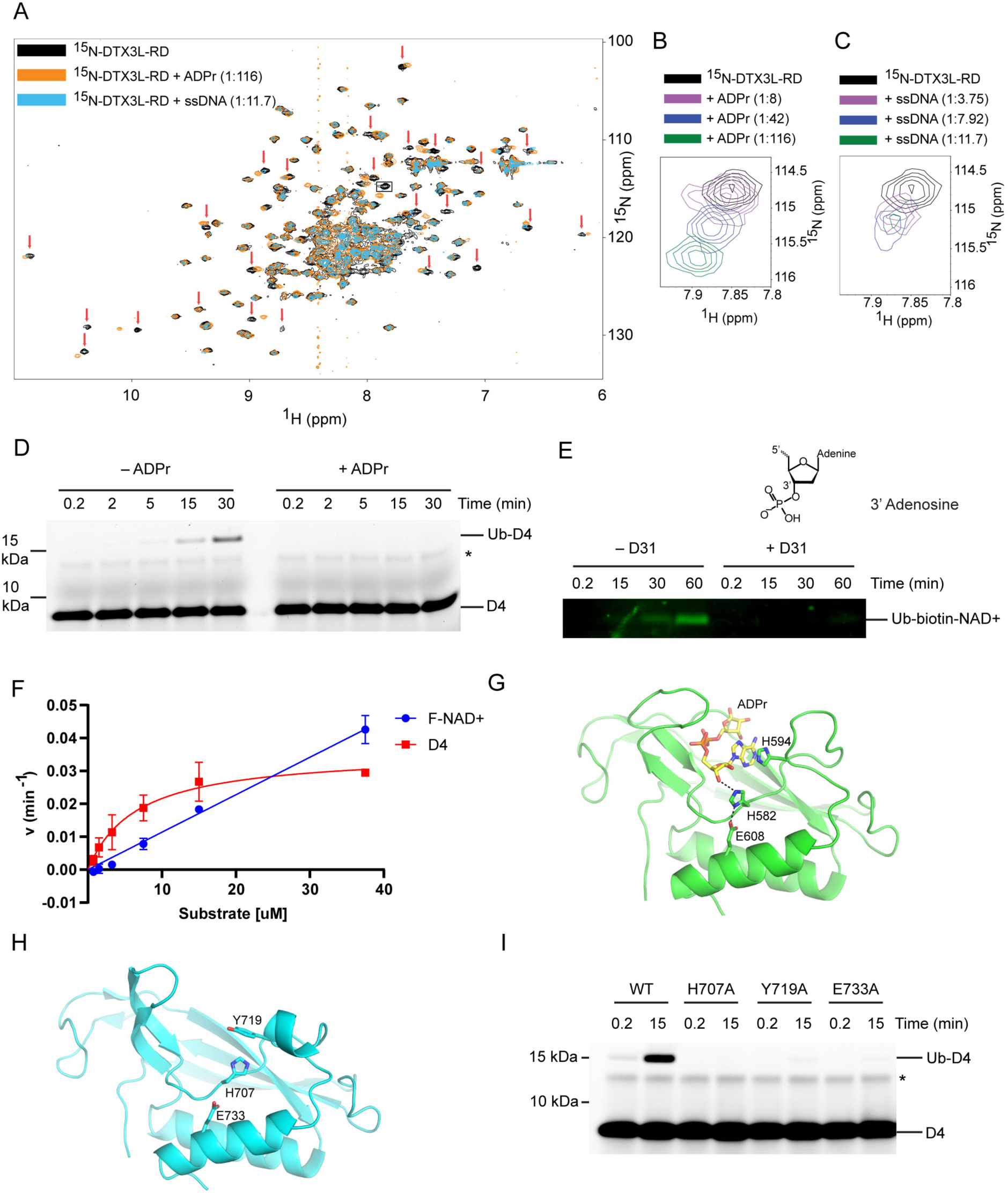
DTX3L DTC domain binds and facilitates Ub-DNA formation. **(A)** ^1^H-^15^N HSQC spectra of ^15^N-DTX3L-RD (black), ADPr-^15^N-DTX3L-RD (orange), and ssDNA D30-^15^N-DTX3L-RD (blue). Red arrows indicate cross peaks that shift upon titrating with ADPr or ssDNA. **(B)** Close-up view of the cross peak indicated by black box in (A) upon titration of specified molar ratios of ADPr with ^15^N-DTX3L-RD. **(C)** Close-up view of the cross peak indicated by black arrow in (A) upon titration of specified molar ratios of ssDNA D30 with ^15^N-DTX3L-RD. **(D)** Fluorescently detected SDS-PAGE gel of *in vitro* ubiquitination of 6-FAM-labelled ssDNA D4 by DTX3L-RD in the presence of E1, UBE2D2, Ub, Mg^2+^-ATP and treated with excess ADPr. **(E)** Western Blot of *in vitro* ubiquitination of biotin-NAD^+^ by DTX3L-RD in the presence of E1, UBE2D2, Ub, Mg^2+^-ATP and treated with excess ssDNA D31. **(F)** Kinetics of Ub-D4 and Ub-F-NAD^+^ formation catalysed by DTX3L-RD. Data from two independent experiments (n=2) were fitted with the Michaelis–Menten equation and *k*_cat_/*K*_m_ value for D4 (5457 M^−1^ min^−1^) calculated. *k*_cat_/*K*_m_ value for F-NAD^+^ (1190 M^−1^ min^−1^) was estimated from the slope of the linear portion of curve. **(G)** Structure of DTX2-DTC domain (green) bound to ADPr (yellow) (PDB: 6Y3J). The sidechains of H582, H594 and E608 are shown in sticks. Hydrogen bonds are indicated by dotted lines. **(H)** Structure of DTX3L-DTC domain (cyan; PDB: 3PG6). The sidechains of H707, Y719 and E733 are shown in sticks. **(I)** Fluorescently detected SDS-PAGE gel of *in vitro* ubiquitination of 6-FAM-labelled ssDNA D4 by full length DTX3L WT, H707A, Y719A and E733A in the presence of E1, UBE2D2, Ub, Mg^2+^-ATP. Asterisks in (D) and (I) indicate contaminant band from ssDNA. Uncropped gel images of (D), (E) and (I) are shown in fig. S5.

To assess DTX3L-RD preference for nucleic acid or ADPr, we performed kinetic analysis of Ub-D4 and Ub-F-NAD^+^ formation by DTX3L-RD using D4 and 6-Fluo-10-NAD^+^ (F-NAD^+^), respectively, as substrates. DTX3L-RD displayed a *k*_cat_ value of 0.0358 ± 0.0034 min^−1^ and a *K*_m_ value of 6.56 ± 1.80 μM for Ub-D4 formation, whereas the Michaelis-Menten curve did not reach saturation for Ub-F-NAD^+^ formation (Fig. 4F and fig. S2, D-G). Comparison of the estimated catalytic efficiency (*k*_cat_/*K*_m_ = 5457 M^−1^ min^−1^ for D4 and estimated *k*_cat_/*K*_m_ = 1190 M^−1^ min^−1^ for F-NAD^+^; Fig. 4F) suggested that DTX3L-RD exhibited 4.5-fold higher catalytic efficiency for D4 than F-NAD^+^. This difference primarily results from a better *K*_m_ value for D4 compared to F-NAD^+^. Although DTX3L-RD showed weak *K*_m_ for F-NAD^+^, it displays a higher rate for converting F-NAD^+^ to Ub-F-NAD^+^ at higher substrate concentration (Fig. 4F). Thus, substrate concentration will play a role in determining the preference.

Previously, we demonstrated the importance of DTX2 His^594^ located in the DTC pocket for the interaction with ADPr (Fig. 4G) (*10*). We examined the effect of mutating the corresponding residue (Tyr^719^) in DTX3L (Fig. 4H) and found that the Y719A mutant was defective in ubiquitination of ssDNA (Fig. 4I). Two catalytic residues in the DTC domain of DTX2(*9*), His^582^ and Glu^608^, have been previously identified as important for the ubiquitination of ADPr, and are conserved across the DTX family members (Fig. 4, G and H and fig. S3A). Based on the similarities in mechanism and conservation of the binding site between ADPr and ssDNA, we next investigated if these catalytic residues were involved in the ubiquitination of ssDNA. Substitution of His^707^ by alanine in FL DTX3L completely abolished the formation of Ub-DNA, whilst substitution of Glu^733^ by alanine considerably impaired product formation (Fig. 4I). To test whether this mechanism was conserved across ssNAs, we repeated the mutant DTX3L ubiquitination assay with ssRNA R4 and found similar results (fig. S2H). Overall, the data support the idea that DTX3L utilises the ADPr-binding pocket and the same catalytic residues in the DTC domain to bind and catalyse ubiquitination of ssNAs.

### Ubiquitination of ssDNA is unique to DTX3 and DTX3L

Since the DTC domains are conserved between DTX family members and share the ability to ubiquitinate ADPr (*7*), we tested whether ubiquitination of ssDNA was a universal feature of the DTX family. Unexpectedly, only DTX3L and to a lesser extent DTX3 were able to ubiquitinate ssDNA under our assay conditions (Fig. 5, A–D). Several potential explanations were considered for this preference: (1) other DTX DTCs might not bind ssDNA; (2) other DTXs might favour different sequences of ssDNA; (3) or the arrangement of the RING and DTC domains in relation to each other might play a crucial role in catalysis. To probe this further, we titrated DTX3L-RING with excess DTX3L-DTC and tracked the formation of Ub-DNA, comparing the ability of the separate domains to ubiquitinate DNA *in trans* with that of DTX3L-RD. No ubiquitination of ssDNA was observed even at a 10:1 molar ratio of DTC to RING domain (Fig. 5E), indicating that the ubiquitination of ssDNA only occurs at an appreciable rate *in cis*. Next, we examined whether DTX2 could catalyse the ubiquitination of ssDNA using 9 different ssDNA sequences and found that DTX2 was unable to ubiquitinate them (Fig. 5F). Lastly, we produced domain-swapped variants of DTX2 and DTX3L in which their respective RING and DTC domains were swapped (DTX2^R^-DTX3L^D^ and DTX3L^R^-DTX2^D^) and examined the ability of these chimeras to ubiquitinate biotin-NAD^+^ or ssDNA D4. We hypothesised that the DTX3L DTC domain of DTX2^R^-DTX3L^D^ would be sufficient to enable binding and ubiquitination of ssDNA. Both chimeras were competent in performing ubiquitination of biotin-NAD^+^ (Fig. 5G) but were unable to produce Ub-DNA (Fig. 5H). These results suggest that the specific arrangement of the RING and DTC domains in DTX3L is crucial in specifying Ub modification of ssDNA. Taken together, we propose a mechanism for the ubiquitination of ssNAs by the DTX3L-RD domains (Fig. 5I): the DTC domain binds ssNAs, the RING domain recruits E2∼Ub, and the RD domains bring the thioester linked Ub into proximity with the 3’ end of ssNA. This allows the catalytic residues in the DTC domain to enable the covalent attachment of Ub to the 3’ hydroxyl group on the 3’-end of ssNA. Subsequently, the complex dissociates, releasing ssNA-Ub. The substrates for the DTC domain also include ADPr, NAD^+^, and ADPr-modified nucleic acids and proteins.

**Figure 5.**
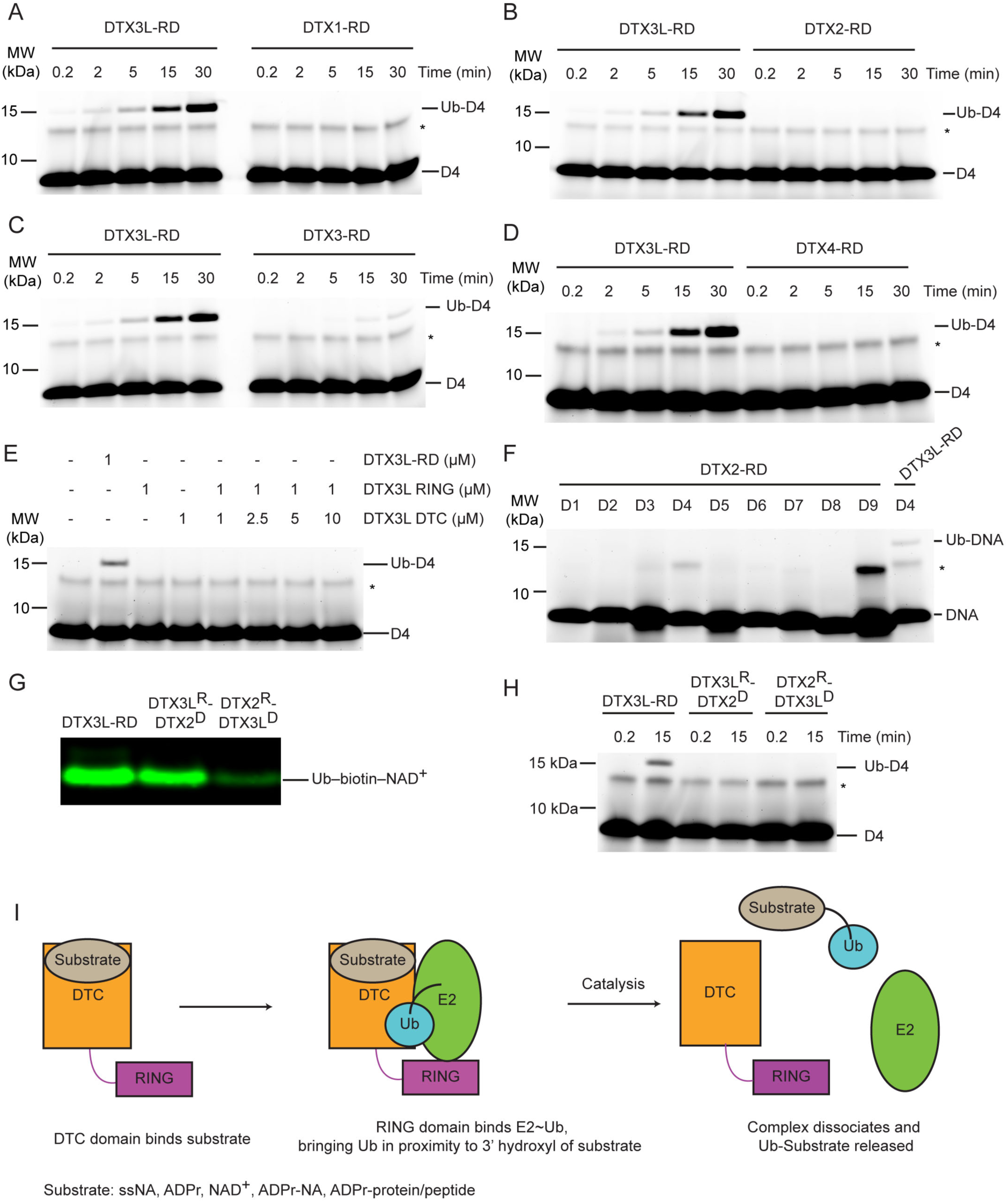
Select DTX RING-DTC domains catalyse ubiquitination of ssDNA. **(A)** Fluorescently detected SDS-PAGE gel of *in vitro* ubiquitination of 6-FAM-labelled ssDNA D4 by DTX3L-RD or DTX1-RD in the presence of E1, UBE2D2, Ub, Mg^2+^-ATP. **(B)** As in (A) but with DTX2-RD. **(C)** As in (A) but with DTX3-RD. **(D)** As in (A) but with DTX4-RD. **(E)** As in (A) but with DTX3L-RD or DTX3L RING with increasing concentrations of DTX3L DTC. **(F)** Fluorescently detected SDS-PAGE gel of *in vitro* ubiquitination of 6-FAM-labelled ssDNA D1-9 by DTX2-RD. A reaction with DTX3L-RD and 6-FAM-labelled ssDNA D4 was included as a positive control. **(G)** Western blot of *in vitro* ubiquitination of biotin-NAD^+^ in the presence of E1, UBE2D2, Ub, Mg^2+^-ATP, NAD^+^, biotin-NAD^+^ with either DTX3L-RD, DTX3L^R^-DTX2^D^, or DTX2^R^-DTX3L^D^ and separated by SDS-PAGE. **(H)** Fluorescently detected SDS-PAGE gel of *in vitro* ubiquitination of 6-FAM-labelled ssDNA D4 by DTX3L-RD, DTX3L^R^-DTX2^D^, or DTX2^R^-DTX3L^D^ in the presence of E1, UBE2D2, Ub, Mg^2+^-ATP. **(I)** Schematic diagrams showing the proposed mechanism of ubiquitination of substrates by DTX3L-RD. Asterisks in (A–F) and (H) indicate contaminant band from ssDNA. Uncropped gel images of (A–H) are shown in fig. S5.

## DISCUSSION

The diversity of non-proteinaceous substrates that have been demonstrated to undergo ubiquitination has been steadily expanding over the last few years (*5, 7, 21*). Here, we report additional examples of non-canonical substrates, namely ssRNA, ssDNA and dsDNA with a 3’ overhang of at least two nucleotides, that undergo ubiquitination by DTX3L. Our biochemical and NMR analyses show that the DTX3L DTC domain binds ssNAs, whilst the RING domain is required for formation of the Ub-NA product. Mutational studies herein reveal three residues, His^707^, Tyr^719^ and Glu^733^, within the DTC domain that are required for the formation of Ub-NA. These residues are conserved within the DTC domain, are known to coordinate ADPr (*7, 10*) and are proposed to constitute a catalytic triad essential for DTX E3-mediated ubiquitination of ADPr (*9*). These findings, coupled with knowledge that the 3’-hydroxyl of the nucleotide ribose is the modification site, suggest a potential similarity in the mechanism of Ub modification of ssDNA, ssRNA, and ADPr by DTX3L.

The chemical structures of ssNAs and ADPr share a ribose-5-phosphate moiety and potentially an adenine base, depending on the ssNA sequence. While our data suggest that DTX3L prefers to ubiquitinate ssDNA ending with adenosine and to some extent guanosine – both purine bases, pyrimidine bases are also tolerated. This raises uncertainty about how DTX3L DTC binds ssNA. Our experiments with dsDNA with a 3’ overhang in one strand showed that dsDNA with a single nucleotide overhang cannot be ubiquitinated, whereas dsDNA with a two-nucleotide overhang was ubiquitinated. Notably, alterations in the sequence of the last two nucleotides at the 3’ end influenced Ub-DNA product formation (Fig. 3, A and B). These observations suggest that the DTX3L DTC NA-binding pocket may accommodate a dinucleotide. Upon examining the ssDNA sequences utilised in our study (Fig. 1F and 3A; Table 1), it appears that the DTX3L DTC pocket might accommodate diverse sequences in the last two nucleotides at the 3’ end, with adenine being the preferred last nucleotide and cytosine being the least favoured. Despite the sequence conservation of the RING-DTC domain with other DTX proteins, our biochemical data revealed that only DTX3L and DTX3 were able to ubiquitinate ssDNA. A comparison of the aligned sequences of the DTX RING-DTC domains reveals some differences (fig. S3A). The RING domains of DTX3L and DTX3 lack insertions found in DTX1, DTX2 and DTX4, making their RING domains appear smaller (fig. S3, A–C). Furthermore, DTX3L and DTX3 lack an AR motif found near the DTC pocket. The absence of the AR motif causes a slight conformational change near the DTC pocket, resulting in an extended β-strand and the loss of a bulge containing the arginine residue (fig. S3, A, D, E). Consequently, DTX3L has an extended groove adjacent to the DTC pocket that could accommodate one or more of the nucleotides 5’ to the targeted terminal nucleotide. Replacing DTX2-RD’s DTC domain with that of DTX3L produced a chimeric construct that was able to ubiquitinate NAD^+^ but not ssDNA (Fig. 5, G and H), suggesting that additional factors beyond the catalytic residues and ssDNA binding properties of the DTC domain likely play a role in ubiquitination. Our findings show that this reaction only occurs *in cis*, pointing to the orientation of the RING relative to the DTC domain as an important feature in the formation of Ub-DNA. Future structural characterisation of DTX3L-RD binding to ssDNA or ssRNA should provide insights into its binding mode and sequence selectivity, thereby potentially revealing whether the biological substrate is ssDNA, ssRNA or both.

Typically, KH domains contain a GXXG motif within the loop between the first and second α helix (*22*). However, analysis of the sequence of the KHL domains in DTX3L shows these domains lack this motif. Multiple studies have shown that mutation in this motif abolishes binding to nucleic acids (*23–26*). Our findings show the DTX3L DTC domain binds nucleic acids but whether the KHL domains contribute to nucleic acid binding requires further investigation. Additionally, the structure of the first KHL domain was recently reported and shown to form a tetrameric assembly (*20*). Our analysis with DTX3L 232-C, which lacks the first KHL domain and RRM, indicate that it can still bind ssDNA and ssRNA. Despite this, a more detailed analysis will be required to determine whether oligomerization plays a role in nucleic acid binding and ubiquitination.

In recent years, several studies have unveiled diverse reactions catalysed by DTX3L. It forms a heterodimeric complex with PARP9, facilitating the direct ubiquitination of substrates such as H2B (*27*). This complex can recruit PARylated substrates, either through PARP9 macrodomains (*27*) or DTX3L DTC domain (*10*), for direct substrate ubiquitination. Notably, it can catalyse the transfer of Ub to NAD^+^ or ADPr (*7–9*), including ADPr moieties of PARylated substrates, encompassing both proteins and nucleic acids (*9*). Moreover, we have expanded its substrate repertoire by demonstrating its ability to directly ubiquitinate the 3’ end of ssNAs. Whether this reaction occurs in cells and what the native substrate (RNA, DNA or both) are yet to be determined. Due to the labile nature of the ribose ester bond and cleavage of Ub conjugates by DUBs, as well as the digestion of nucleic acids by nucleases, our ability to probe the Ub modification of ssNAs in a cellular context has so far been limited. Connecting these chemical reactions to precise biological functions requires carefully designed experiments aimed at distinguishing these functions. Based on the known functions of the DTX3L/PARP9 complex and the findings of this study, we propose several hypotheses for future research. The DTX3L/PARP9 complex has a known involvement in DNA damage repair, utilising PARP9’s macrodomains to recognise PAR and assist in complex recruitment to the damaged foci (*8*). RNA itself contributes to DNA damage repair (*28*). Likewise, these damage sites may involve DNA breaks with 3’ overhangs. Our findings reveal that DTX3L can recognise and ubiquitinate ssRNA, or dsDNA with a 3’ overhang of two or more nucleotides. This observation suggests a possible substrate for DTX3L in DNA damage repair pathways, which requires further investigation. In eukaryotic cells, messenger RNA (mRNA) is a source of ssRNA that commonly possesses a poly-A tail at the 3’ end. Our data indicated that DTX3L can ubiquitinate ssRNA containing a poly-A sequence (Fig. 1G, lane R9), offering a possibility for future exploration. Viruses contain a diverse range of genetic materials including ssNAs, which might be plausible ubiquitination targets. Studies have shown that expression of DTX3L/PARP9 complex is induced after viral infection, and the complex plays a role in antiviral defense mechanisms by ubiquitinating both host and viral proteins (*27, 29*). Ubiquitination of viral ssNAs by DTX3L presents another intriguing hypothesis for future research.

Our study reports the first example of direct ubiquitination of ssNAs by DTX3L and illustrates the possibility of additional roles for ubiquitination beyond that of a conventional post-translational modification.

## Materials And Methods

### Construct generation

Constructs were generated by standard polymerase chain reaction–ligation techniques and verified by automated sequencing. GST-tagged constructs were cloned into a modified form of pGEX4T-3 (GE Healthcare) with a TEV cleavage site and a second ribosomal binding and multiple cloning site (pABLO TEV) and His-tagged constructs were cloned into a modified form of pRSF_Duet-1 (Novagen) with a TEV cleavage site following the hexahistidine tag. All proteins are from human sequences. RD domains are comprised of residues 388-C of DTX1, residues 390-C of DTX2, residues 148-C of DTX3, residues 387-C of DTX4 and residues 544-C of DTX3L. DTC of DTX3L comprises residues 607-C. PARG comprises residues 448-C. USP2 comprises residues 260-C. The complex, DTX3L(232-C)/PARP9(509-C), was cloned into a bicistronic vector in which PARP9(509-C) was cloned into the first MCS of pRSF_Duet-1 with an N-terminal His-MBP tag followed by a TEV cleavage site, and DTX3L(232-C) was cloned into the second MCS untagged. DTX3L 232-C and PARP9 509-C were sufficient to form complex (*27*). DTX3L FL and DTX3L 232-C were cloned into pABLO vector.

### Protein expression and purification

Expression of recombinant proteins was conducted in *Escherichia coli* BL21(DE3) Gold or Rosetta2(DE3) pLysS cells. Cultures were grown in LB broth (Miller) at 37 °C until an OD_600_ of 0.6-0.8 was reached at which point expression was induced with 0.2 mM isopropyl β-D-1-thiogalactopyranoside (IPTG) at 20 °C for 12-16 hours. ^15^N-labelled DTX3L-RD was obtained using M9 minimal medium. Briefly, 20 mL starter cultures were grown in LB medium overnight before cells were pelleted and washed in M9 medium. Each pellet was added to 1L of M9 medium supplemented with: 1 g of ^15^NH_4_Cl, 5 g glucose, 50 mg kanamycin, 1X Vitamin Stock (Gibco), 1 mg D-Biotin and trace metals. Cells were grown until an OD_600_ of 0.6-0.8 was reached at which point expression was induced with IPTG at 20 °C for 20 hours.

Cells were harvested by centrifugation following expression and lysed by microfluidizer or sonicator. Cells expressing GST-tagged proteins were resuspended in 50 mM Tris-HCl (pH 7.6), 200 mM NaCl and 1 mM DTT. Cells expressing His-tagged proteins were resuspended in 25 mM Tris-HCl (pH 7.6), 200 mM NaCl, 15 mM imidazole, 5 mM beta-mercaptoethanol. Cell lysates were cleared by high-speed ultracentrifugation. Clarified lysates were applied to glutathione affinity or Ni^2+^-agarose by incubating for 1 to 2 hours on a rotary shaker at 4°C or using a gravity column. Beads were washed in buffers similar to lysis buffer. ^15^N-DTX3L-RD was eluted with 6xHis intact in 25 mM Tris-HCl (pH 7.6), 200 mM NaCl, 5 mM BME, and 200 mM imidazole, then further purified by ion exchange chromatography, followed by size exclusion chromatography on a Superdex 75 column (GE Healthcare) into buffer containing 20 mM sodium phosphate, (pH 7.0), 100 mM NaCl, 0.02% NaN_3_ and 1 mM TCEP before snap-freezing in liquid nitrogen and storing at −80 °C.

For removal of the GST tag, protein samples were dialysed against 25 mM Tris-HCl (pH 7.6), 150 mM NaCl, and 5 mM BME overnight at 4 °C in the presence of TEV protease or alternatively cleaved on the beads with TEV from a glutathione agarose column in 50 mM Tris-HCl (pH 8.0), 200 mM NaCl, and 5 mM DTT. The dialysed and cleaved samples were loaded onto the same resin and flow through collected to separate from the affinity tags or the remaining uncleaved proteins. Additional purification was performed by size exclusion chromatography on a Superdex 75 column or Superdex 200 column (GE Healthcare), depending on protein size, into 25 mM Tris-HCl (pH 7.6), 150 mM NaCl, and 1 mM DTT before snap-freezing in liquid nitrogen and storing at −80 °C. Protein concentrations were determined using Bio-RAD protein assay.

DTX3L (232-C)/PARP9 (509-C) complex was obtained by expressing the bicistronic construct of His-MBP-PARP9(509-C)/DTX3L(232-C). The complex was purified by Ni^2+^-agarose, cleaved by TEV treatment, loaded back on Ni^2+^ for tag and protease removal, and further purified by Source S cation exchange chromatography and Superdex 200.

Additional proteins: *Arabidopsis thaliana* Uba1 (*30, 31*), UBE2D2 (*30, 32*), Ub (*30, 33*), PARG (*7*) and USP2 (*34*) were purified as in previously described protocols.

### Oligonucleotides

Oligonucleotides were obtained from Integrated DNA Technologies and are listed in Table 1.

### Fluorescence polarisation assay

50 nM 6-FAM-labelled nucleic acids (Integrated DNA Technologies) were incubated with 0-5 µM proteins in reaction buffer (20 mM Hepes pH 7.0, 1 mM MgCl_2_, 50 mM NaCl, filter sterilised). Fluorescence polarisation measurements were taken using Tecan Spark (λex=485 nm, λem=535 nm). Fold change was calculated based on the change in polarisation compared to the fluorescent ligand in the absence of protein. Data were plotted in Prism (GraphPad).

### DNA and RNA ubiquitination assays

UBA1 (0.2 µM), UBE2D2 (5 µM) and Ub (50 µM) were incubated in 50 mM Tris-HCl, 5 mM MgCl_2_, 50 mM NaCl and 5 mM ATP at room temperature for 15 minutes to allow for generation of E2∼Ub. DTX (1 µM) and nucleic acid (3 µM) were incubated together at room temperature for 10 minutes prior to mixing with other components. After components were combined, reactions were incubated at 37 °C. Aliquots were taken immediately (indicated by 0.2 min time point) and at specified subsequent time points. The reactions were halted by addition of 2X Loading Dye containing 250 mM DTT and resolved by SDS-PAGE (NuPAGE 4-12% Bis-Tris, Invitrogen). For FAM-labelled nucleic acids, gels were visualised using Typhoon FLA 700 with λex=473 nm and Y520 emission filter (Cytiva Life Sciences). For IRDye 800 labelled nucleic acids, gels were scanned with an Odyssey CLx Imaging System (LI-COR Biosciences). For Fig. 1J and 5, A, B, C and D, 25 µM D4 was used. For Fig. 1L, 25 µM D11 was used. For assays with USP2, Benzonase and PARG, reactions were conducted as above and after UBA1 (0.2 µM), UBE2D2 (5 µM), Ub (50 µM), DTX3L-RD (2 µM) and D4 or R4 (2 µM) were incubated together for 30 minutes, reactions were quenched with 250 mM DTT and buffer exchanged into 25 mM Tris-HCl (pH 7.6), 150 mM NaCl, 5 mM MgCl_2_ and 1 mM DTT using a 7 kDa Zeba Spin column. USP2 (1.35 µM), Benzonase (1 µM) or PARG (1 µM) was added and incubated for a further 30 minutes at 37 °C. Subsequently, an aliquot was mixed with 2X Loading Dye and resolved by SDS-PAGE. For assays conducted at pH 9.5, reactions were conducted as above and after UBA1 (0.2 µM), UBE2D2 (5 µM), Ub (50 µM), DTX3L-RD (1 µM) and D4 (3 µM) were incubated together for 30 minutes, reactions were quenched with 250 mM DTT and buffer exchanged into 25 mM Tris-HCl (pH 9.5), 150 mM NaCl and 1 mM DTT using a 7 kDa Zeba Spin column. Then 2 mM EDTA was added, and the reaction was incubated at 37 °C. Aliquots were taken at indicated time points, mixed with 2X Loading Dye, and resolved by SDS-PAGE. For assays conducted with NH_2_OH treatment, reactions were conducted as above and after UBA1 (0.2 µM), UBE2D2 (5 µM), Ub (50 µM), DTX3L-RD (1 µM) and D4 (3 µM) were incubated together for 30 minutes. Reactions were quenched with 250 mM DTT and buffer exchanged into 25 mM Tris-HCl (pH 9), 150 mM NaCl and 1 mM DTT using a 7 kDa Zeba Spin column. Then 1.5 M NH_2_OH was added, and the reaction was incubated at 37 °C. Aliquots were taken at indicated time points, mixed with 2X Loading Dye, and resolved by SDS-PAGE.

### Autoubiquitination assay

UBA1 (0.2 µM), UBE2D2 (5 µM) and fluorescently-labelled Ub (*35*) (25 µM) were incubated in 50 mM Tris-HCl, 5 mM MgCl_2_, 50 mM NaCl and 5 mM ATP at room temperature for 15 minutes to allow for generation of E2∼Ub. DTX3L (1 µM) was added and reactions were incubated at 20 °C. Aliquots were taken at indicated time points, reactions halted by addition of 2X Loading Dye and resolved by SDS-PAGE. Gels were scanned with an Odyssey CLx Imaging System.

### Lysine discharge assay

UBA1 (0.5 µM), UBE2D2 (25 µM) and Ub (25 µM) were incubated in 50 mM Tris-HCl, 5 mM MgCl_2_, 50 mM NaCl and 5 mM ATP at room temperature for 15 minutes to allow for generation of E2∼Ub. Charged reactions were stopped with 30 mM EDTA and T0 sample taken. DTX3L (1 µM) and lysine (300 mM) were added, and reactions were incubated at room temperature. Aliquots were taken at indicated time points, reactions halted by addition of 2X Loading Dye and resolved by SDS-PAGE. Gels were scanned with a ChemiDoc (BioRad).

### Competition ubiquitination assays

UBA1 (0.2 µM), UBE2D2 (5 µM) and Ub (50 µM) were incubated in 50 mM Tris-HCl, 5 mM MgCl_2_, 50 mM NaCl and 5 mM ATP at room temperature for 15 minutes to allow for generation of E2∼Ub. DTX3L-RD (1 µM) and D4 (3 µM) were incubated together at room temperature for 10 minutes prior to mixing with other components. After components were combined, 1 mM ADPr was added and reactions were incubated at 37 °C. Aliquots were taken at indicated time points, reactions were halted by addition of 2X Loading Dye containing 250 mM DTT and resolved by SDS-PAGE. Gels were visualised using Typhoon FLA 700 with λex=473 nm and Y520 emission filter. For Fig. 4E, DTX3L-RD (1 µM) and biotin-NAD^+^ (5 µM) were incubated together at room temperature for 10 minutes prior to mixing with other components. After components were combined, 2 mM D31 was added and reactions were incubated at 30 °C. Aliquots were taken at indicated time points, reactions were halted by addition of 2X Loading Dye containing 250 mM DTT and resolved by SDS-PAGE. The samples were transferred to nitrocellulose membrane and blocked in 5% BSA. Membranes were incubated with DyLight 800 conjugated NeutrAvidin (Thermo Fisher Scientific, cat. no. 22853; 1:10,000) and visualised using an Odyssey CLx Imaging System.

### Kinetic analysis of Ub-D4 and Ub-F-NAD+ formation

UBA1 (0.2 µM), UBE2D2 (5 µM) and Ub (50 µM) were incubated in 50 mM Tris-HCl, 5 mM MgCl_2_, 50 mM NaCl and 5 mM ATP at room temperature for 15 minutes to allow for generation of E2∼Ub. DTX (1 µM) and D4 or F-NAD^+^ (Biolog Life Science Institute, cat. no. N 023) at varying concentrations (0-37.5 µM) were incubated together at room temperature for 10 minutes prior to mixing with other components. Aliquots were taken after 15 minutes, reactions were halted by addition of 2X Loading Dye containing 250 mM DTT and resolved by SDS-PAGE. Gels were visualised using Typhoon FLA 700 with λex=473 nm and Y520 emission filter. A known amount of D4 was visualised on the gel alongside D4 ubiquitination reactions. A known amount of F-NAD^+^ was pipetted onto Whatman filter paper and scanned alongside gel of F-NAD^+^ ubiquitination reactions. Ub-D4 and Ub-F-NAD^+^ were quantified and initial rates expressed as nmoles of product formed per minute per moles of DTX3L-RD. Data from two independent experiments were fitted to the Michaelis-Menten equation using Prism. The Michaelis-Menten curve for Ub-F-NAD^+^ was not saturated therefore *k*_cat_*/K*_m_ was estimated from the slope of the linear portion of the curve.

### Biotin-NAD^+^ ubiquitination assay

UBA1 (0.2 µM), UBE2D2 (5 µM), and Ub (50 µM) were incubated in 50 mM HEPES-NaOH, pH 7.5, 5 mM MgCl_2_, 50 mM NaCl and 5 mM ATP at room temperature for 10 minutes to allow for generation of E2∼Ub. DTX (1 µM), NAD^+^ (200 µM) and biotin-NAD^+^ (10 µM) were added and reactions were incubated at 20 °C for one hour. Reactions were halted by addition of 3X Loading Dye containing 250 mM DTT and resolved by SDS-PAGE. The samples were transferred to nitrocellulose membrane and blocked in 5% BSA. Membranes were incubated with DyLight 800 conjugated NeutrAvidin (Thermo Fisher Scientific, cat. no. 22853; 1:10,000) and visualised using an Odyssey CLx Imaging System.

### Solution NMR experiments

All NMR data were acquired using a Bruker Avance III 600-MHz spectrometer equipped with a cryogenic triple resonance inverse probe. DTX3L-RD samples were prepared at ∼100 μM in 20 mM sodium phosphate, (pH 7.0), 100 mM NaCl, 0.02% NaN_3_, 1 mM TCEP and 5% D_2_O with 0.00025% TSP. Experiments were carried out at 298 K, and ^1^H-^15^N HSQC spectra were recorded with 16 scans using 200 complex points with a sweep width of 36 parts per million (ppm) in the ^15^N dimension. All spectra were processed with 256 points in the indirect dimension using Bruker TopSpin 3.5 and analysed using CcpNmr AnalysisAssign (*36*).

## Acknowledgements

We thank Core Services and Advanced Technologies at the Cancer Research UK Scotland Institute (C596/A17196 and A31287) with particular thanks to Molecular Technology services; Catherine Winchester for her assistance in critically reviewing this manuscript.

## Author contributions

E.L.D., C.C., L.B., S.F.A., and D.T.H. generated constructs and purified proteins. E.L.D. and C.C. carried out the investigation. E.L.D. and B.O.S. performed NMR and analysed the data. C.C. and T.S. performed ssRNA and ssDNA binding assays and analysed with M.B. E.L.D. and D.T.H. wrote the manuscript. All authors commented on the manuscript.

## Data and materials availability

Original and uncropped images are presented in Supplemental Figures 4 and 5. Materials are available from the corresponding author (D.T.H.) upon request.

---

**Figure S1.**
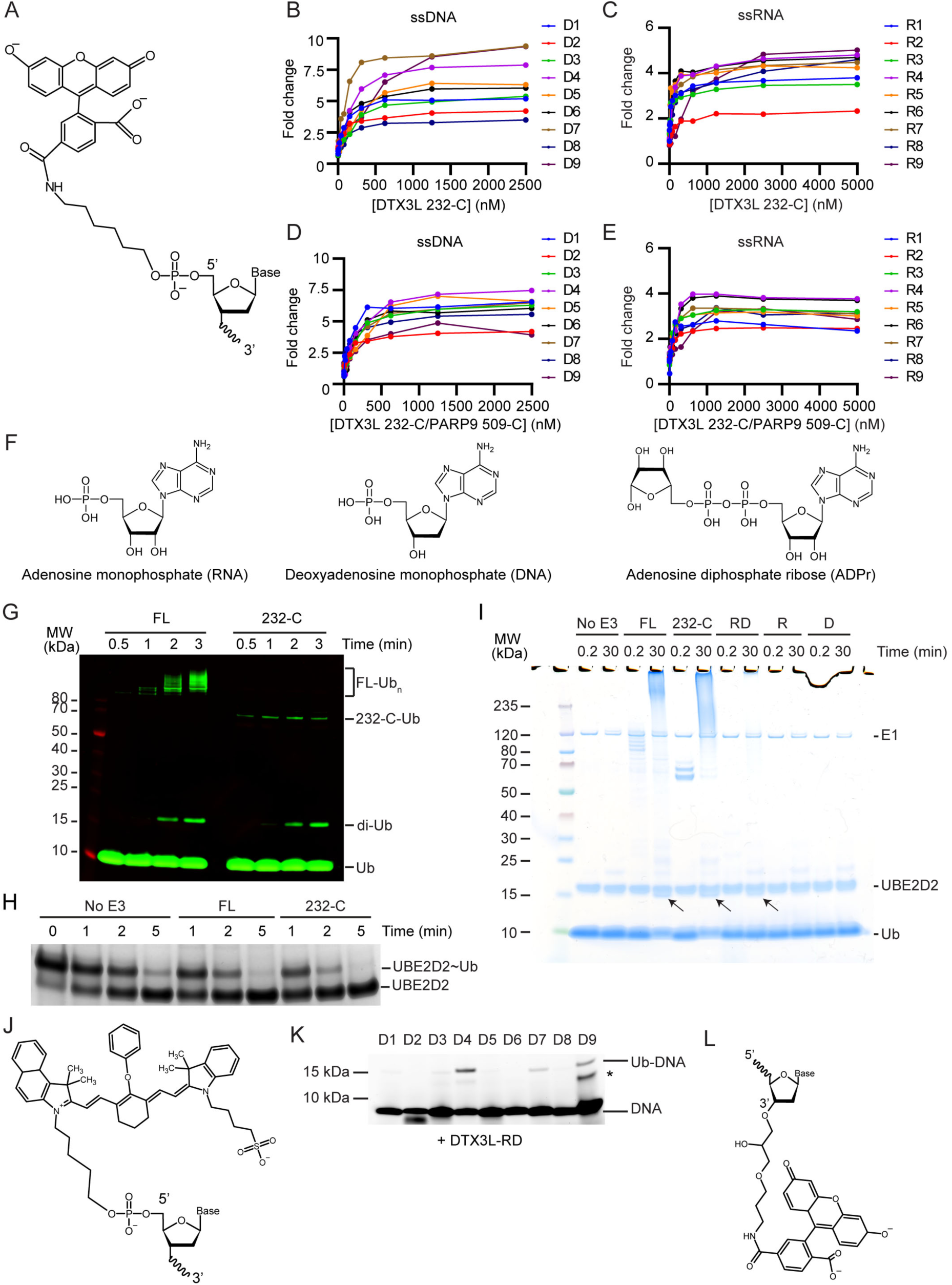
DTX3L binds and ubiquitinates ssNAs. **(A)** Schematic of the 5’ 6-FAM modification. **(B)** Fold change of fluorescence polarisation of 6-FAM-labelled ssDNA D1-9 oligonucleotides upon titrating with DTX3L 232-C. **(C)** As in (B) but with 6-FAM-labelled ssRNA R1-9 oligonucleotides. **(D)** As in (B) but with DTX3L 232-C/PARP9 509-C. **(E)** As in (D) but with 6-FAM-labelled ssRNA R1-9. **(F)** Schematic of RNA, DNA and ADPr. **(G)** Fluorescently detected SDS-PAGE gel of *in vitro* autoubiquitination of DTX3L FL and 232-C in the presence of E1, UBE2D2, fluorescent-labelled Ub and Mg^2+^-ATP. **(H)** Coomassie stained SDS-PAGE gel of lysine discharge reactions showing disappearance of UBE2D2∼Ub over time in the presence of FL, 232-C or absence of E3. **(I)** Coomassie stained SDS-PAGE gel of *in vitro* ubiquitination of 6-FAM-labelled ssDNA D4 by DTX3L variants in the presence of E1, UBE2D2, Ub, Mg^2+^-ATP (from Fig. 1H). Arrows indicate potential ∼15 kDa Ub-DNA band. **(J)** Schematic of IRDye® 800 modification. **(K)** Fluorescently detected SDS-PAGE gel of *in vitro* ubiquitination of 6-FAM-labelled ssDNA D1-9 in the presence of E1, UBE2D2, Ub, Mg^2+^-ATP by DTX3L-RD. **(L)** Schematic of the 3’ 6-FAM modification. Asterisks in (K) indicate contaminant band from ssDNA. Uncropped gel images of (H) and (K) are shown in fig. S5.

**Figure S2.**
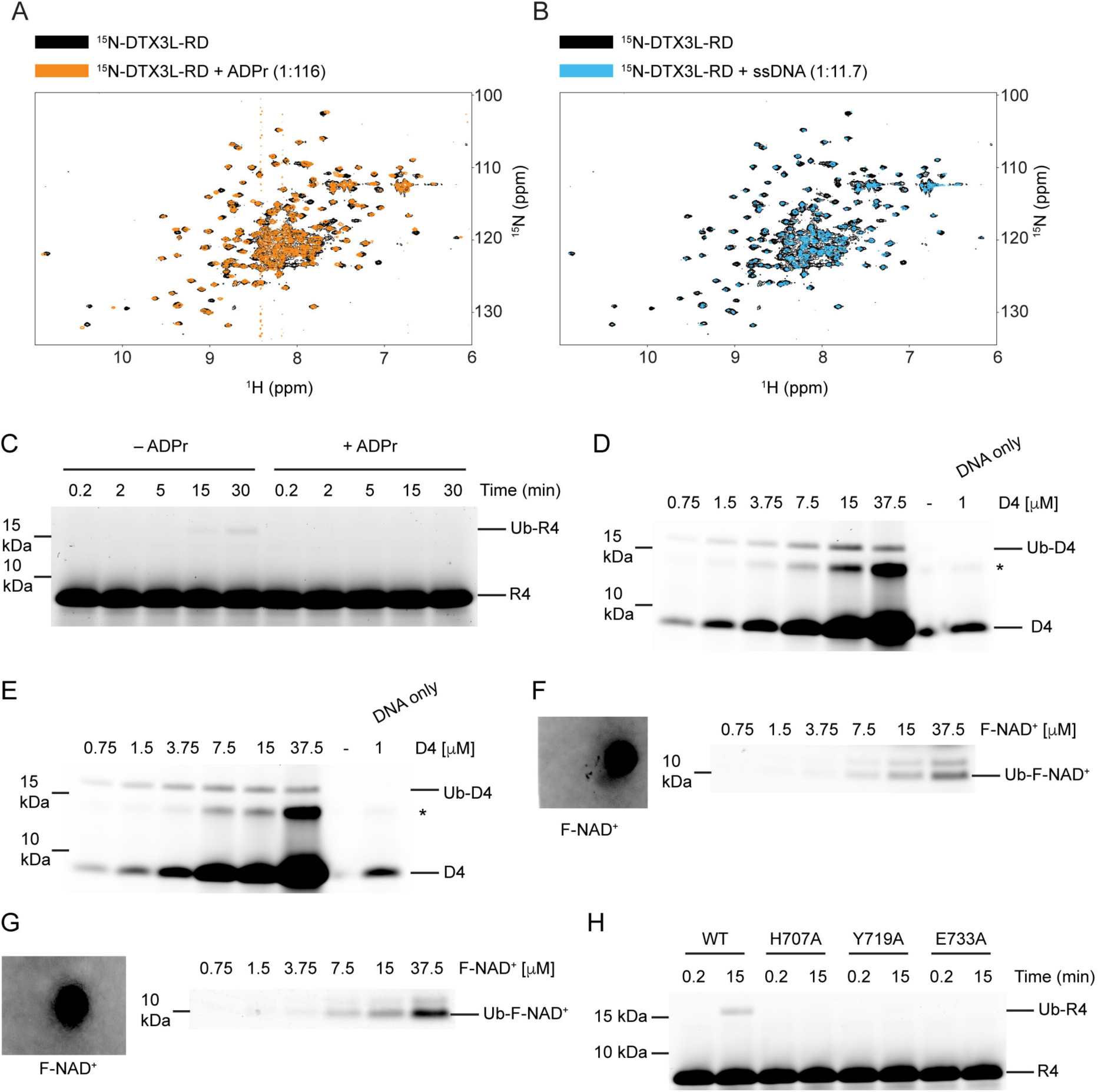
DTX3L-RD binds ADPr and ssNA. **(A)** ^1^H-^15^N HSQC spectra of ^15^N-DTX3L-RD (black) and after the addition of ADPr (orange) (related to Fig. 4A). **(B)** ^1^H-^15^N HSQC spectra of ^15^N-DTX3L-RD (black) and after the addition of ssDNA D30 (blue) (related to Fig. 4A). **(C)** Fluorescently detected SDS-PAGE gel of *in vitro* ubiquitination of 6-FAM-labelled ssRNA R4 by DTX3L-RD in the presence of E1, UBE2D2, Ub, Mg^2+^-ATP and treated with excess ADPr. **(D)** Fluorescently detected SDS-PAGE gel of *in vitro* ubiquitination of increasing concentrations of 6-FAM-labelled ssDNA D4 by DTX3L-RD in the presence of E1, UBE2D2, Ub, Mg^2+^-ATP. **(E)** Replicate of (D). **(F)** Fluorescently detected SDS-PAGE gel of *in vitro* ubiquitination of increasing concentrations of F-NAD^+^ by DTX3L-RD in the presence of E1, UBE2D2, Ub, Mg^2+^-ATP. A known volume of 100 µM F-NAD^+^ was pipetted onto Whatman filter paper and scanned alongside the gel for quantification. **(G)** Replicate of (F). **(H)** Fluorescently detected SDS-PAGE gel of *in vitro* ubiquitination of 6-FAM-labelled ssRNA R4 by FL DTX3L WT, H707A, Y719A and E733A in the presence of E1, UBE2D2, Ub, Mg^2+^-ATP. Asterisks in (D) and (E) indicate contaminant band from ssDNA. Uncropped gel images of (C-H) are shown in fig. S5.

**Figure S3.**
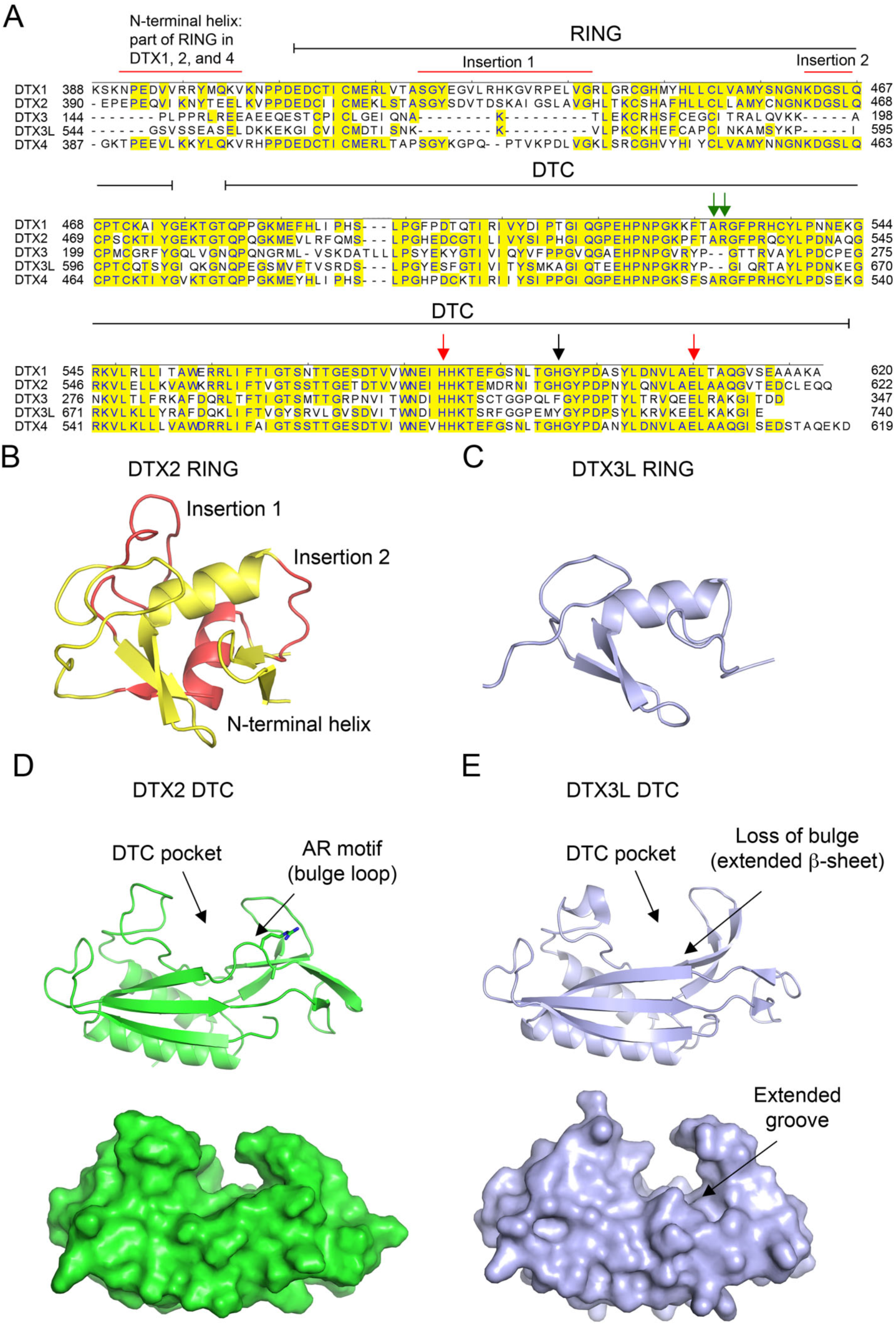
Properties of DTX family DTC domains. (**A**) Sequence alignment of human DTX family RING-DTC domains. Starting and ending residues of RING-DTC domains are indicated. Conserved residues are highlighted in yellow. A black arrow indicates residue that stacks with ADPr, red arrows indicate conserved catalytic residues, and green arrows indicate AR motif present in DTX1, 2, and 4. Insertions are indicated. (**B**) Structure of DTX2 RING domain (from PDB: 6Y3J). The insertions and N-terminal helix are colored in red and the conserved RING domain region is colored yellow. (**C**) Alphafold2 model of DTX3L RING domain (light blue) displayed in the same orientation as in B. (**D**) Top: cartoon representation of structure of DTX2 DTC domain (green; PDB: 6Y3J). Sidechains of AR motif are shown as sticks. The DTC pocket that binds ADPr and the bulged AR motif are indicated by arrows. Bottom: as in top panel but with surface representation. (**E**) Top: cartoon representation of structure of DTX3L DTC domain (light blue; PDB: 3PG6). Bottom: as in top panel but with surface representation. The absence of an AR motif in DTX3L causes a slight structural change near the ADPr-binding pocket, leading to the loss of the bulged loop, which results in a slightly extended groove.

**Figure S4.**
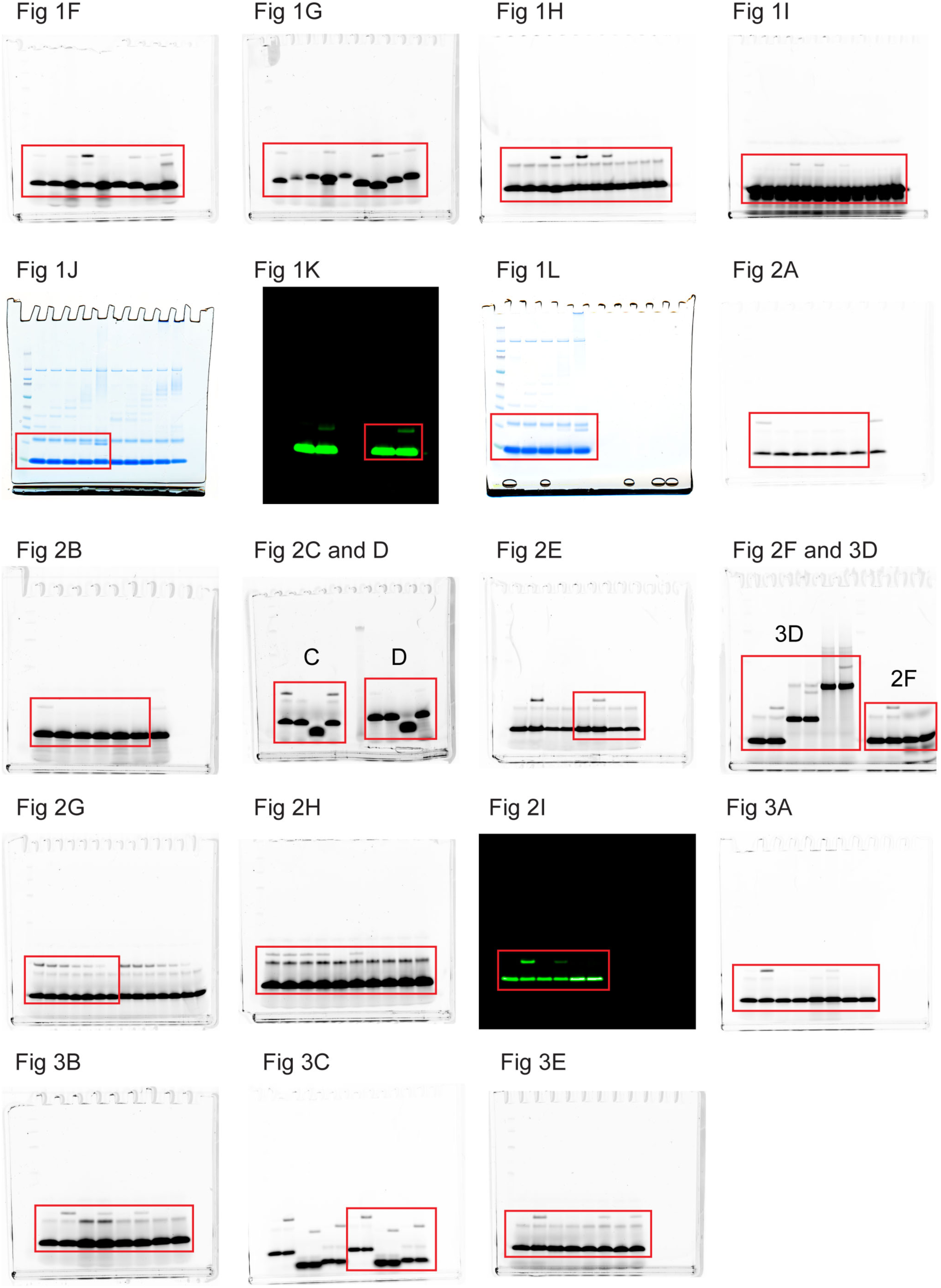
Full images of gels from Figure 1-3. Red boxes denote where images were cropped.

**Figure S5.**
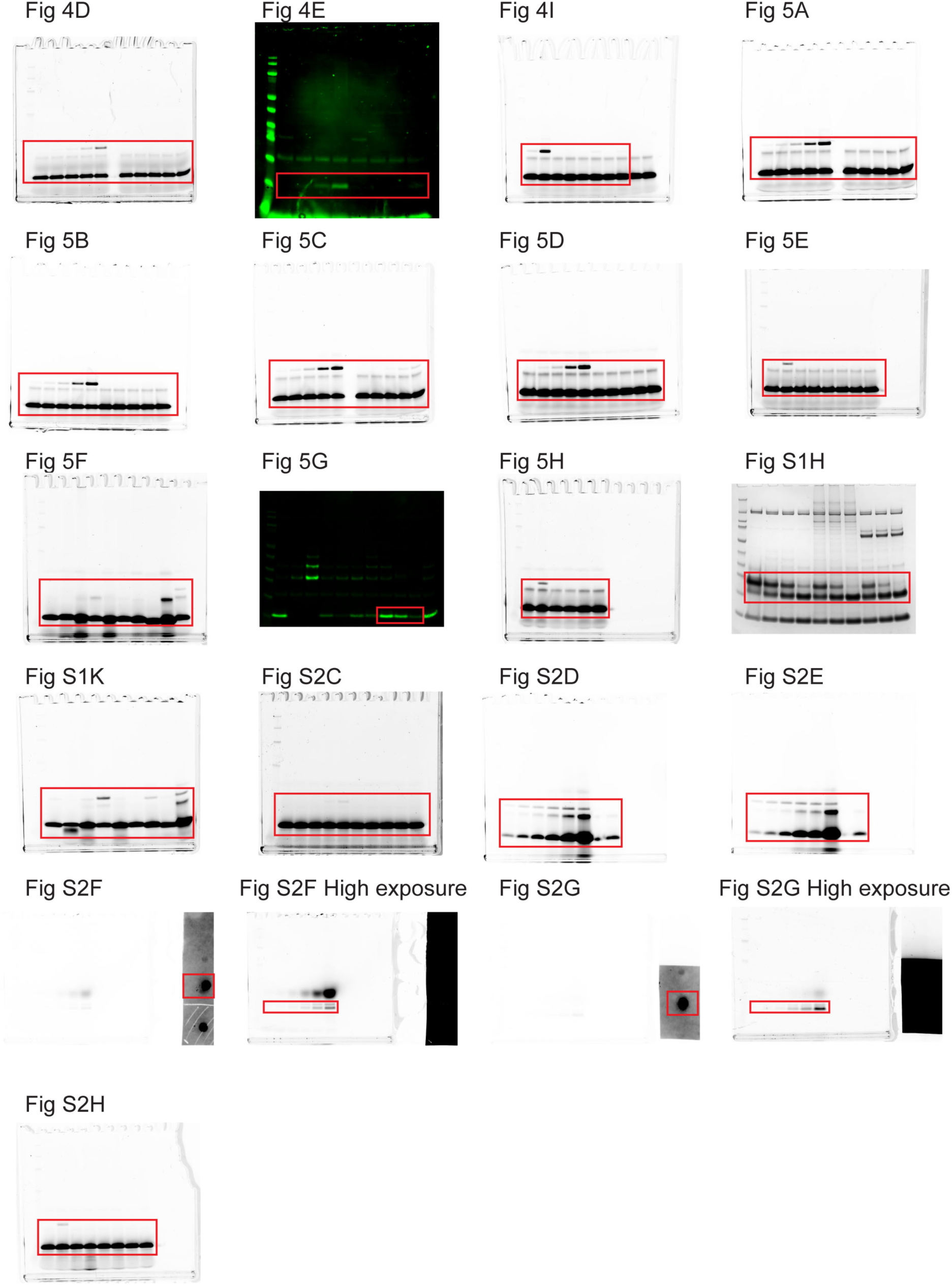
Full images of gels from Figure 4-5 and S1-2. Red boxes denote where images were cropped.

## References

1. R. Yau, M. Rape, The increasing complexity of the ubiquitin code. Nat. Cell Biol. 18, 579–586 (2016).

2. X. Wang, R. A. Herr, M. Rabelink, R. C. Hoeben, E. J. H. J. Wiertz, T. H. Hansen, Ube2j2 ubiquitinates hydroxylated amino acids on ER-associated degradation substrates. J. Cell Biol. 187, 655–668 (2009).

3. Y. Shimizu, Y. Okuda-Shimizu, L. M. Hendershot, Ubiquitylation of an ERAD Substrate Occurs on Multiple Types of Amino Acids. Mol. Cell 40, 917–926 (2010).

4. X. Wang, R. A. Herr, W. J. Chua, L. Lybarger, E. J. Wiertz, T. H. Hansen, Ubiquitination of serine, threonine, or lysine residues on the cytoplasmic tail can induce ERAD of MHC-I by viral E3 ligase mK3. J. Cell Biol. 177, 613–624 (2007).

5. E. G. Otten et al., Ubiquitylation of lipopolysaccharide by RNF213 during bacterial infection. Nature 594, 111–116 (2021).

6. I. R. Kelsall et al., HOIL-1 ubiquitin ligase activity targets unbranched glucosaccharides and is required to prevent polyglucosan accumulation. EMBO J. 41, e109700 (2022).

7. C. Chatrin et al., Structural insights into ADP-ribosylation of ubiquitin by Deltex family E3 ubiquitin ligases. Sci. Adv. 6, eabc0418 (2020).

8. C. S. Yang et al., Ubiquitin Modification by the E3 Ligase/ADP-ribosyltransferase Dtx3L/Parp9. Mol. Cell 66, 503–503 (2017).

9. K. Zhu et al., DELTEX E3 ligases ubiquitylate ADP-ribosyl modification on protein substrates. Sci. Adv. 8, eadd4253 (2022).

10. S. F. Ahmed et al., DELTEX2 C-terminal domain recognizes and recruits ADP-ribosylated proteins for ubiquitination. Sci. Adv. 6, eabc0629 (2020).

11. R. J. Deshaies, C. A. Joazeiro, RING domain E3 ubiquitin ligases. Annu. Rev. Biochem. 78, 399–434 (2009).

12. K. Takeyama et al., The BAL-binding Protein BBAP and Related Deltex Family Members Exhibit Ubiquitin-Protein Isopeptide Ligase Activity. J. Biol. Chem. 278, 21930–21937 (2003).

13. S. Vyas et al., Family-wide analysis of poly(ADP-ribose) polymerase activity. Nat. Commun. 5, (2014).

14. K. Zhu et al., DELTEX E3 ligases ubiquitylate ADP-ribosyl modification on nucleic acids. Nucleic Acids Res. 52, 801–815 (2024).

15. J. Jumper et al., Highly accurate protein structure prediction with AlphaFold. Nature 596, 583–589 (2021).

16. J. L. Baber, D. Libutti, D. Levens, N. Tjandra, High Precision Solution Structure of the C-terminal KH Domain of Heterogeneous Nuclear Ribonucleoprotein K, a c-myc Transcription Factor. J. Mol. Biol. 289, 949–962 (1999).

17. J. L. Baber, D. Levens, D. Libutti, N. Tjandra, Chemical Shift Mapped DNA-Binding Sites and 15N Relaxation Analysis of the C-Terminal KH Domain of Heterogeneous Nuclear Ribonucleoprotein K. Biochemistry 39, 6022–6032 (2000).

18. A. Cléry, M. Blatter, F. H. T. Allain, RNA recognition motifs: boring? Not quite. Curr. Opin. Struct. Biol. 18, 290–298 (2008).

19. Y. Ashok et al., Reconstitution of the DTX3L–PARP9 complex reveals determinants for high-affinity heterodimerization and multimeric assembly. Biochem. J. 479, 289–304 (2022).

20. C. Vela-Rodríguez et al., Oligomerization mediated by the D2 domain of DTX3L is critical for DTX3L-PARP9 reading function of mono-ADP-ribosylated androgen receptor. Protein Sci. 33, e4945 (2024).

21. I. R. Kelsall, J. Zhang, A. Knebel, J. S. C. Arthur, P. Cohen, The E3 ligase HOIL-1 catalyses ester bond formation between ubiquitin and components of the Myddosome in mammalian cells. Proc. Natl. Acad. Sci. U.S.A. 116, 13293–13298 (2019).

22. N. V. Grishin, KH domain: one motif, two folds. Nucleic Acids Res. 29, 638–643 (2001).

23. D. Hollingworth et al., KH domains with impaired nucleic acid binding as a tool for functional analysis. Nucleic Acids Res. 40, 6873–6886 (2012).

24. T. Talwar et al., The DEAD-box protein DDX43 (HAGE) is a dual RNA-DNA helicase and has a K-homology domain required for full nucleic acid unwinding activity. J. Biol. Chem. 292, 10429–10443 (2017).

25. M. Yadav et al., The KH domain facilitates the substrate specificity and unwinding processivity of DDX43 helicase. J. Biol. Chem. 296, 100085 (2021).

26. C. L. Lin, Y. T. Wang, W. Z. Yang, Y. Y. Hsiao, H. S. Yuan, Crystal structure of human polynucleotide phosphorylase: insights into its domain function in RNA binding and degradation. Nucleic Acids Res. 40, 4146–4157 (2012).

27. Y. Zhang et al., PARP9-DTX3L ubiquitin ligase targets host histone H2BJ and viral 3C protease to enhance interferon signaling and control viral infection. Nat. Immunol. 16, 1215–1227 (2015).

28. A. S. Bader, B. R. Hawley, A. Wilczynska, M. Bushell, The roles of RNA in DNA double-strand break repair. Br. J. Cancer 122, 613–623 (2020).

29. J. Huang et al., DTX3L Enhances Type I Interferon Antiviral Response by Promoting the Ubiquitination and Phosphorylation of TBK1. J. Virol. 97, e0068723 (2023).

30. M. Gabrielsen et al., A General Strategy for Discovery of Inhibitors and Activators of RING and U-box E3 Ligases with Ubiquitin Variants. Mol. Cell 68, 456–470.e410 (2017).

31. H. Dou, L. Buetow, G. J. Sibbet, K. Cameron, D. T. Huang, BIRC7–E2 ubiquitin conjugate structure reveals the mechanism of ubiquitin transfer by a RING dimer. Nat. Struct. Mol. Biol. 19, 876–883 (2012).

32. H. Dou, L. Buetow, A. Hock, G. J. Sibbet, K. H. Vousden, D. T. Huang, Structural basis for autoinhibition and phosphorylation-dependent activation of c-Cbl. Nat. Struct. Mol. Biol. 19, 184–192 (2012).

33. S. Volk, M. Wang, C. M. Pickart, Chemical and genetic strategies for manipulating polyubiquitin chain structure. Methods Enzymol. 399, 3–20 (2005).

34. M. Renatus et al., Structural Basis of Ubiquitin Recognition by the Deubiquitinating Protease USP2. Structure 14, 1293–1302 (2006).

35. H. M. Magnussen et al., Structural basis for DNA damage-induced phosphoregulation of MDM2 RING domain. Nat. Commun. 11, 2094 (2020).

36. S. P. Skinner, R. H. Fogh, W. Boucher, T. J. Ragan, L. G. Mureddu, G. W. Vuister, CcpNmr AnalysisAssign: a flexible platform for integrated NMR analysis. J. Biomol. NMR 66, 111–124 (2016).

